# Genetic and phenotypic consequences of local transitions between sexual and parthenogenetic reproduction in the wild

**DOI:** 10.1101/2022.11.02.514965

**Authors:** Soleille Morelli Miller, Katarina C Stuart, Nathan William Burke, Lee Ann Rollins, Russell Bonduriansky

## Abstract

Transitions from sexual to asexual reproduction have occurred in numerous lineages across the tree of life, but it remains unclear why asexual populations rarely persist. In facultatively parthenogenetic animals, all-female populations can arise when males are absent or become extinct, and such populations can give rise to obligately asexual species. Facultative parthenogens could therefore shed light on the initial stages of transitions to asexuality, and the factors that determine the success or failure of asexual populations. Here, we describe a novel spatial mosaic of mixed-sex and all-female populations of the facultatively parthenogenetic Australian phasmid *Megacrania batesii*, and use this system to investigate the consequences of reproductive mode variation in the wild. Analysis of single nucleotide polymorphisms (SNPs) indicated multiple, independent transitions between reproductive modes. As expected, all-female populations had drastically reduced heterozygosity and genetic diversity relative to mixed-sex populations. However, we found few consistent differences in fitness-related traits between population types. All-female populations exhibited more frequent and severe (non-functional) wing deformities, but did not show higher rates of appendage loss. All-female populations also harbored more parasites, but only in certain habitats. Reproductive mode explained little variation in female body size, fecundity, or egg hatch-rate. Our results confirm that transitions to parthenogenetic reproduction can lead to dramatic reductions in genetic diversity and heterozygosity. However, our findings also suggest that asexual *M. batesii* populations consist of high-fitness genotypes that might be able to thrive for many generations, perhaps until they encounter a drastic environmental change to which they are unable to adapt.

## Introduction

The pervasiveness of sexual reproduction in animals has remained a mystery for over a century. Parthenogenetic strategies avoid physically costly sexual encounters, enjoy increased colonialization ability, and do not spend resources creating males that contribute minimally to offspring (Maynard Smith, 1978). Yet sex is nearly ubiquitous across the animal kingdom.

Efforts to explain the prevalence of sexual strategies have sought to identify the advantages that sexual reproduction offers, particularly in the long term (reviewed in Barton & Charlesworth, 1998; Otto & Gerstein, 2006; Hadany & Comeron, 2008; Neiman & Shwander, 2011). Sex can be considered beneficial because it generates genotypic diversity, thereby potentially facilitating adaptation (Fisher-Muller effect: Fisher, 1930; Muller, 1932; Crow & Kimura, 1965; Park and Krug, 2013; Cooper, 2007; Becks & Agrawal, 2012; Colegrave, 2002). Alternative hypotheses focus instead on the long-term consequences of asexuality (Bell, 1982; Agrawal, 2006; Howard & Lively, 1994; Park, Jokela & Michalakis, 2010). In particular, the accumulation of deleterious alleles in non-recombining animals could result in a reduction of mean fitness due to increased mutation load (i.e., Mullers ratchet, Muller 1964) which can lead to population extinction (Lynch & Gabriel, 1990; Lynch et. al., 1993; Chao, 1990; Loewe & Cutter, 2008). These theories focus on the consequences of reproducing asexually in the long-term but, to explain the prevalence of sexual reproduction in nature, it is also important to understand what happens in the short-term, during the initial stages of transitions to asexuality. In particular, what are the factors that often enable sexual populations to outcompete or prevent invasion by asexual strategies?

In animals, asexual lineages always arise from sexual ancestors (Butlin, 2002) but the process of transition from sexual to asexual reproduction and the short-term consequences of such transitions are not well understood, especially across natural populations. A common approach is to compare costs and benefits of sex and parthenogenesis between closely related sexual and asexual species (Bast et. al., 2018; Kimmerer 1994; Tarkhnishvili et. al., 2010). Consequently, most knowledge of shifts to asexuality comes from studying ancient transitions (e.g., Loewe & Lamatsch, 2008; Schlupp, 2010; Fontaneto, 2012; Larose, Parker & Schwander, 2018). While such an approach has yielded important insights into how asexual lineages may have persisted following ancient transitions to parthenogenesis, it cannot reveal the factors and processes involved in the early stages of transitions to asexual reproduction. Comparing related species also introduces potential confounding factors since genuine consequences of reproducing asexually cannot be differentiated from genetic changes associated with speciation.

Facultatively parthenogenetic animals, in which every female can engage in sexual or asexual reproduction depending on whether mating and fertilization occurs (Normark, 2003), provide an opportunity to disentangle species-specific factors from direct consequences of asexuality. Facultatively parthenogenic animals can reap the benefits of both modes of reproduction while avoiding many of the costs (D’Souza & Michiels, 2010). Facultative parthenogenesis can also be an intermediate stage between obligate sexuality and obligate asexuality (Schwander et. al., 2010), and so studying the genetic and phenotypic differences between sexually and asexually reproducing populations within the same species of a facultative parthenogen could help reveal factors that promote or impede the long-term persistence of asexual populations, and the potential for transitions to obligate asexuality. Many facultatively parthenogenetic animals exhibit a geographic pattern where some populations reproduce sexually while others reproduce asexually. This phenomenon, known as geographic parthenogenesis (GP), has been reported in several taxa, including Opiliones (Burns, Hedin & Tsurusaki, 2018), mayflies (Tojo, Sekiné & Matsumoto, 2006), phasmids (Law & Crespi, 2002; Morgan-Richards, Trewick & Stringer, 2010), and others (reviewed in Glesener & Tilman, 1978). The geographic patterns found in GP animals provide an opportunity to investigate the immediate effects that follow the establishment of asexual populations, a process that can involve unmated females colonizing new areas, or males becoming locally extinct. The immediate genetic and phenotypic effects of such transitions could determine whether asexual populations persist or suffer rapid extinction. Yet, very little is known about the relative performance of females in all-female (asexual) versus mixed-sex (sexual) populations of such species.

The Australian Peppermint Stick Insect, *Megacrania batesii*, is a facultatively parthenogenic species endemic to the wet tropics of far-north Queensland, Australia (Cermak & Hasenpusch, 2000), as well as the Solomon Islands and other Pacific islands (Van Herwaarden, 1998). When *M. batesii* females mate, they create a mixed-sex brood of sexually produced offspring. However, females can also reproduce via thelytoky, whereby females that avoid mating produce all-female broods from unfertilized eggs. While some phasmids can produce male offspring from unfertilized eggs (Pijnacker & Ferwerda, 1980; Scali, 2009; Brock et. al., 2018), it is currently unknown how common parthenogenetically produced males are in *M. batesii*. Cermak & Hasenpusch (2000) reported the existence of two isolated all-female populations at the southern end of the Australian range of *M. batesii*, but pilot surveys identified a complex geographic mosaic of populations with contrasting sex ratios throughout the species range. Females from all-female locations do not show obvious morphological differentiation from females found in mixed-sex locations and experimental population crosses in our laboratory (to be reported elsewhere) showed that these populations are inter-fertile. Since sexual reproduction is believed to be the ancestral reproductive mode in phasmids (Schwander & Crespi, 2009), we assume that all-female populations of this species are descended from mixed-sex, sexual populations.

Our first aim was to describe geographic variation in sex ratio and reproductive mode in wild populations of *M. batesii*. Our second aim was to use these populations to investigate the consequences of transitions to asexual reproduction in wild populations within their natural habitats. In particular, we aimed to determine whether incipient transitions to asexuality (i.e., parthenogenetic reproduction) are associated with reduced genetic diversity and heterozygosity, as predicted by theory and shown by empirical studies on other species (Stenberg & Saura, 2009). We also wanted to determine whether such genetic changes are associated with reduced phenotypic performance in fitness-related traits, including female body size, fecundity, egg hatching success, appendage deformity and loss, and parasite load. A number of studies have compared the performance of sexual and asexual animals in the laboratory (Browne et. al., 1988; Kenny 1996; Cullum 1997; Mee et. al., 2011; Sukumaran & Grant, 2013). However, given the strong environment-dependence of life history and fitness (Saether & Engen 2015), there is a need for research on wild populations in fully natural environments. *M. batesii* occurs in two types of habitat: rainforest along beach margins (where it feeds on *Pandanus* sp. host-plants), and swamp within closed-canopy rainforest (where it feeds on *Benstonea* sp. host-plants). We therefore also asked whether phenotypic and genotypic parameters are affected by habitat type. Because we are interested in how reproductive mode transitions affect functionality in natural environments, we report phenotypic data collected from individuals observed in natural populations, rather than from their lab-reared descendants.

## Methods

### Study locations

We investigated a spatial mosaic of populations with either mixed sex or all-female sex ratios along ~ 33km of coastline between Forest Creek and Emmagen Creek. We also surveyed two isolated all-female locations (reported previously by Cermak & Hasenpusch, 2000) located ~ 160-180 km south of this area at Etty Bay and Bingil Bay (see Fig. 4a). *M. batesii* were observed and sampled from habitat along Cape Tribulation Road and Kimberley Road, in Forest Creek Village, Cow Bay Village, Cape Tribulation Village and adjacent beaches, at Emmagen Creek, and at Etty Bay and Bingil Bay, in Queensland, Australia.

### Population sex ratio and phenotyping

A total of 1,324 wild *M. batesii* individuals were photographed on their host plants at 25 locations in 2019 (N = 258), 2020 (N = 283), 2021 (N = 259), and 2022 (N = 524) to quantify sex ratio and collect phenotypic data. At each location, we scanned host plants (*Pandanus* sp. or *Benstonea* sp.) and recorded the presence, developmental status (nymph or adult), sex of adults and some nymphs that were developed enough to distinguish sex, and pairing status (single female, single male or male-female pair) of each *M. batesii* individual seen. Given the very effective crypsis of *M. batesii*, it is likely that we failed to spot some individuals at each location, but our surveys should nonetheless represent differences between locations in the relative abundance, sex ratio and phenotype of *M. batesii* individuals. The *M. batesii* sites surveyed were classified as “all-female” if only females were seen, and “mixed-sex” if both sexes were seen (see Fig. 3a). Localities with numerous individuals could be differentiated as “all-female” or “mixed-sex” because males, when present, are easy to distinguish from females by their body shape and wing length (see Fig. 1). In mixed-sex sites, many adult females are guarded by males (Boldbaatar et al. 2022). One site where a few females (but no males) were found was designated a “putative all-female” location (B1). One site (NN) was consistently all-female for 2019-2021 but, in 2022, males were found at that site. This site is hereafter referred to as a “transitional” site and was excluded from phenotypic analyses comparing sexual and asexual locations.

**Figure 1:**
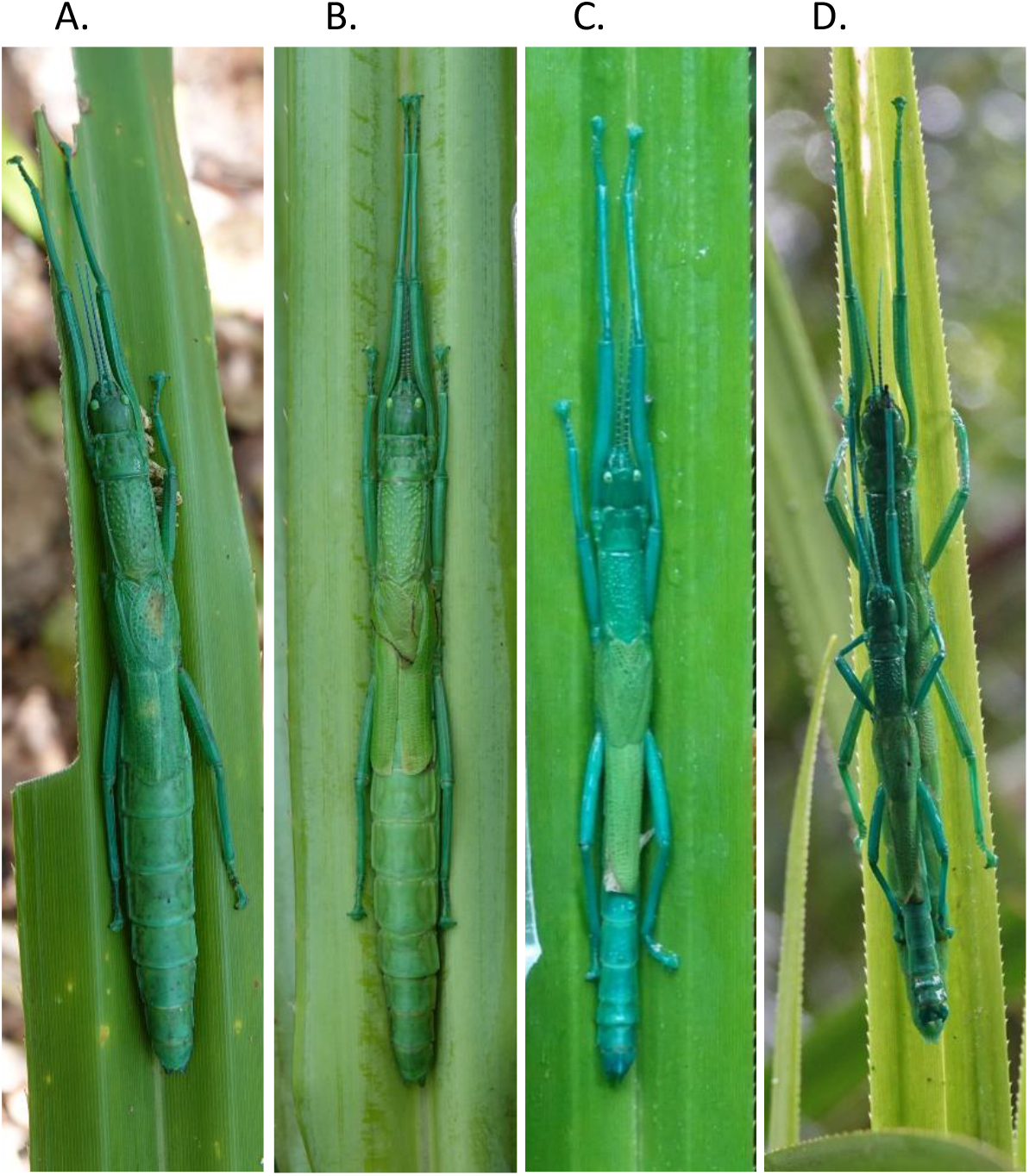
*Megacrania batesii* individuals and pairs: (A) adult female from an all-female site; (B) adult female from a mixed-sex site; (C) adult male; (D) female guarded by male.

Each adult and nymph seen in the field was imaged in situ on its host plant using a Sony RX10-IV camera. When possible, a transparent plastic scale attached to a thin metal rod was held next to the adult insects so that body size measurements could be obtained from the image (Fig. S1). In most cases, this procedure did not appear to disturb the insects since no movement or defensive reaction (retreating, jumping off the leaf, or spraying of defensive fluid) was observed. From the images, ImageJ (Schneider, Rasband & Eliceiri, 2012) was used to measure the length of the pro-thoractic notum (protonum) and scale. Pronotum length was then calculated in cm as a measure of body size. We measured pronotum length because this part of the body was consistently visible in the photos, including photos of females being guarded by males. The photos were also used to record the number of missing legs and antennae. Leg and antennae loss in arthropods is an indication of poor condition that could result from predator attack or problems with moulting (Maruzzo et. al., 2005). For each individual, we also recorded the presence of black spots indicative of fungal infection, the presence of ectoparasites (mites or biting midges), and the presence and severity of wing deformities. Normal wings are held flat over the dorsal thorax of the insect, with one wing overlapping the other. Mild wing deformities were defined as normally shaped wings held apart (not overlapping), whereas severe wing deformities were defined as wings that were widely separated, partly or entirely missing, or highly abnormal in shape.

In 2019 and 2020, we collected 1,418 eggs from several wild-collected *M. batesii* females and sexed 719 hatchlings from those eggs to verify that females at the all-female locations were reproducing asexually while females at the mixed-sex locations were reproducing sexually. Wild-collected females were kept in mesh cages in for 7-12 days and fed on *Benstonea* sp. or *Pandanus* sp. leaves, and then released at the site of capture. Females that were mating or guarded by males when collected were housed together with the male. Eggs laid by wild-collected females were counted and then kept in containers with moist cocopeat at ~27°C until hatching in controlled-temperature rooms at The University of New South Wales, Sydney. Fecundity was quantified for each wild-collected female as the number of eggs laid per day, and egg viability was quantified as the hatching rate of these eggs. Hatchlings were sexed based on the morphology of the 8^th^ and 9^th^ abdominal sternites (Fig. S2), and hatchling sex ratio was used to assess reproductive mode in the natural source locations. Fertilized eggs hatch into both males and females whereas unfertilized eggs typically hatch into females only. Thus, when only female hatchlings were obtained from eggs laid by wild-collected females, we inferred that that the females at that location reproduced only via parthenogenesis; whereas when male hatchlings were obtained, we inferred that sexual reproduction had taken place. Based on data from laboratory crosses (not shown), spontaneous production of males from unfertilized eggs is impossible or extremely rare (< 1/1000 eggs) in *M. batesii* (unpublished data).

### DNA sampling, extraction and purification

In 2020 and 2021, mid-legs were collected from wild nymphs or adults for DNA sequencing by pinching the leg with sterile tweezers, causing the insect to autotomize the pinched leg. A few additional samples were obtained from adults that died in cages during egg collection, or from whole first-instar nymphs collected in the field. Samples were stored in pure ethanol at 4°C or without ethanol at −80°C until DNA extraction.

Isolation of DNA was conducted using the Gentra Puregene Tissue Kit (4g) from Qiagen according to the manufacturer’s protocol, “DNA Purification from Mouse Tail Tissue Using the Gentra Puregene Mouse Tail Kit,” with slight modifications to suit the study species (www.qiagen.com). Specifically, prior to adding 1.5 μl Puregene Proteinase K to the lysate, the samples were subsampled and placed in 2ml free standing screw-capped tubes. Then, 3 × 5mm glass beads and ~5-10 × 1mm silicon beads were placed in the tubes with the samples. The tubes were placed in a FastPrep-24 (MP Biomedical) homogenizer and run on a 2 × 25-second cycle to break down the sample exoskeleton. Following homogenization, 1.5 μl Puregene Proteinase K was added to the lysate and the rest of the manufacturer’s protocol was followed.

Isolates from a total of 188 *M. batesii* individuals were sent to Diversity Arrays Technology Inc. (Canberra, Australia) for whole-genome reduced representation genotyping using the DArT-seq protocol. Genome complexity reduction was performed using a restriction enzyme double digest of PstI and HpaII. Next-generation sequencing of amplification fragments was conducted on the Illumina Hiseq 2500 (www.illumina.com), producing 1,384,779 single-end reads of raw data across all 188 samples. SNPs were then called using the DArTsoft analytical pipeline (Kilian et. al., 2012).

### Quality control filtering

The DArTsoft pipeline was used to calculate quality parameters, including call rate, reproducibility, and polymorphic information content (PIC). The initial data set provided by DArTseq consisted of 12,977 SNPs across the 188 samples. Due to a high percentage of missing data (>30%) and possible mislabeling, 33 samples were omitted from this analysis (see supplementary file “omitted_samples.xls”). A total of 12,518 polymorphic DArT markers were generated for the retained 155 samples. Before quality control filtering, the mean repeatability was 0.987 and over 50% of SNPs had reproducibility over 99% (see Fig. S4). The mean minor allele frequency by locus was 0.18 but varied across locations with asexual sites having higher levels of minor alleles compared to sexual sites. The mean call rate was 0.95 and 95% of SNPs had a call rate above 75%. The PIC value of SNP markers ranged from 0.0 to 0.5 with a mean of 0.239 and median of 0.228.

DArTSeq™ sequences were filtered using dartR (Gruber et. al., 2018). Various filter parameters were tested to determine suitable filters and SNP sequences matching the following parameters were removed from the dataset: average reproducibility < 95%, call rate < 75%, minor allele frequency (MAF) < 0.05, read depth < 3, and Hamming distance > 0.80. Remaining SNPs were assessed for linkage disequilibrium and all SNPs with a hamming distance less than 0.2 were excluded from further analysis. Quality control filtering removed 5,603 SNPs or 43.17% of SNPs (see Table S5). After quality control, 7,374 high quality SNPs were identified for further analysis.

### Phenotypic analysis

All phenotypic analyses were conducted in R, v. 4.0.2 (R Core Team, 2020). The package “MCMCglmm” was used to model variation in phenotypic traits using a mixed model approach (Hadfield, 2010). For each response variable, reproductive mode (assumed to be sexual for mixed-sex sites and asexual for all-female sites), and habitat type (*Benstonea* swamp or *Pandanus* beach) were modeled as fixed effects, and none were transformed. All models also included year, and a matrix representing genetic structure, as random effects.

Each phenotypic response variable was modeled separately with different specifications depending on deviance information criterion (DIC) (see Table S7). Sample sizes differed for different phenotypic variables (for all summary stats see Table S2). Pronotum length was analysed with a Gaussian model. Presence of fungal infections and ectoparasites were combined into one response variable (“parasites”) showing the presence/absence of fungal infections or ectoparasites and analysed with a categorical (binary) model. The numbers of missing legs and antennae were added together and this sum was analysed as a single response variable. The variable was analysed with a Poisson model because there were no instances of a stick insect with all appendages missing, thus there was no upper bound on the variable. Presence/absence of mild and severe wing deformities were analysed with a categorical (binary) model. Hatching success of eggs collected from wild females was also analysed as a binary matrix of successes and failures. Fecundity (number of eggs laid) was analysed using a Poisson model.

Host plant type is strongly correlated with habitat type, with *Pandanus* sp. host plants typically growing along beach margins, and *Benstonea* sp. host plants growing along margins of streams and swamps in closed-canopy rainforest. We therefore characterized habitat type as “*Pandanus*-beach” or “*Benstonea*-swamp”. All models initially included the interaction between reproductive mode and habitat, but this interaction was dropped if its inclusion increased DIC. However, the main effects of reproductive mode and habitat type were retained in all models (see Table S1). For each phenotypic variable, the matrix representing population-level phylogenetic information (i.e., genetic structure) was used to calculate the broad sense phylogenetic heritability (H^2^) and assess the amount of phylogenetic signal. Phylogenetic information used in the models was collected using the *gl.dist.pop()* function in the R package “dartR” and inverse phylogenetic covariance matrices were created using the function *inverseA()*. Phylogenetic information was retained in all models.

### Population structure and genetic diversity

We investigated the genetic relationships between locations by calculating a dissimilarity matrix using the *bitwise.dist()* function in the R package “poppr” (Kamvar, Tabima, & Grünwald, 2014) based on Hamming distances (see Wang et. al., 2015). This matrix was then used with *pvclust()* to create a dendrogram based on correlation distance between individuals and 10,000 bootstrap iterations were done to determine the level of support for each node (Suzuki & Shimodaira, 2006). Several clustering methods were compared (see supp materials “testing_clustering_methods.html”), all methods yielding similar genetic structure. Pairwise *F_st_* values were calculated using the function *stamppFst()* with 10,000 bootstrap iterations used to obtain confidence limits for each pairwise difference (Pembleton, Cogan, & Forster, 2013). Non-Metric Multidimensional Scaling (NMDS) using *metaMDS()* was then used to visualize the structure among samples as a complement to the cluster analysis (Dixon, 2003). Observed and expected heterozygosities as well as *F_is_* of each locality was obtained by *gl.get.diversity()* (see Table S6). Sites with less than two sampled individuals were excluded from these analyses.

Gene flow between sites was estimated using discriminant analysis of principal components (DAPC). DAPC uses K-means and model selection to determine genetic clusters in lieu of known populations or genetic groups (Jombart, Devillard & Balloux, 2010). Seventeen optimal genetic clusters were identified using the Bayesian Information Criterion (see Fig. S3) and DAPC cross validation using the function *xvalDapc()*. Six discriminant analysis axes were retained based on the α score. Membership probabilities were estimated for each individual for each identified group and *ggplot()* was then used to visualize the admixture proportions of each individual’s genotype.

Allelic diversity, Shannon’s diversity, and heterozygosity were quantified using the function *gl.report.diversity()* as described by Sherwin et. al. (2017), and these values were compared between sites (see Fig. S5). Observed heterozygosity (Ho) values were calculated using the function *gl.report.heterozygosity()* and used to visualize heterozygosity of each sample (see Fig. 5c). Isolation by distance was determined using the function *gl.ibd()*. All functions used are from the package “dartR” (Gruber et al., 2018). The data that support the findings of this study are openly available in dryad at https://doi.org/10.5061/dryad.n02v6wx1g.

## Results

### Sex ratio and reproductive mode

Sex ratio was bimodally distributed, with either only females or an approximately equal number of females and males observed at nearly all locations (see Table 2). With one exception (see below), we did not find any locations where adult males were present but rare (and the absence of such locations is supported by population genomic analysis, described below). In several cases, all-female and mixed-sex sites were found to occur in close proximity, and with no obvious barriers to dispersal. For example, mixed-sex locations B4 and VR are separated by < 2km of contiguous rainforest from all-female locations B1 and CB, and the all-female location NS is separated by < 1km of rainforest from adjacent mixed-sex locations (See Fig. 4). Despite the close proximity of males to some all-female sites, sex ratios were consistent over four years of surveys at nearly all locations (see Supplementary file “sex_ratio_by_year.csv”).

**Table 2:**
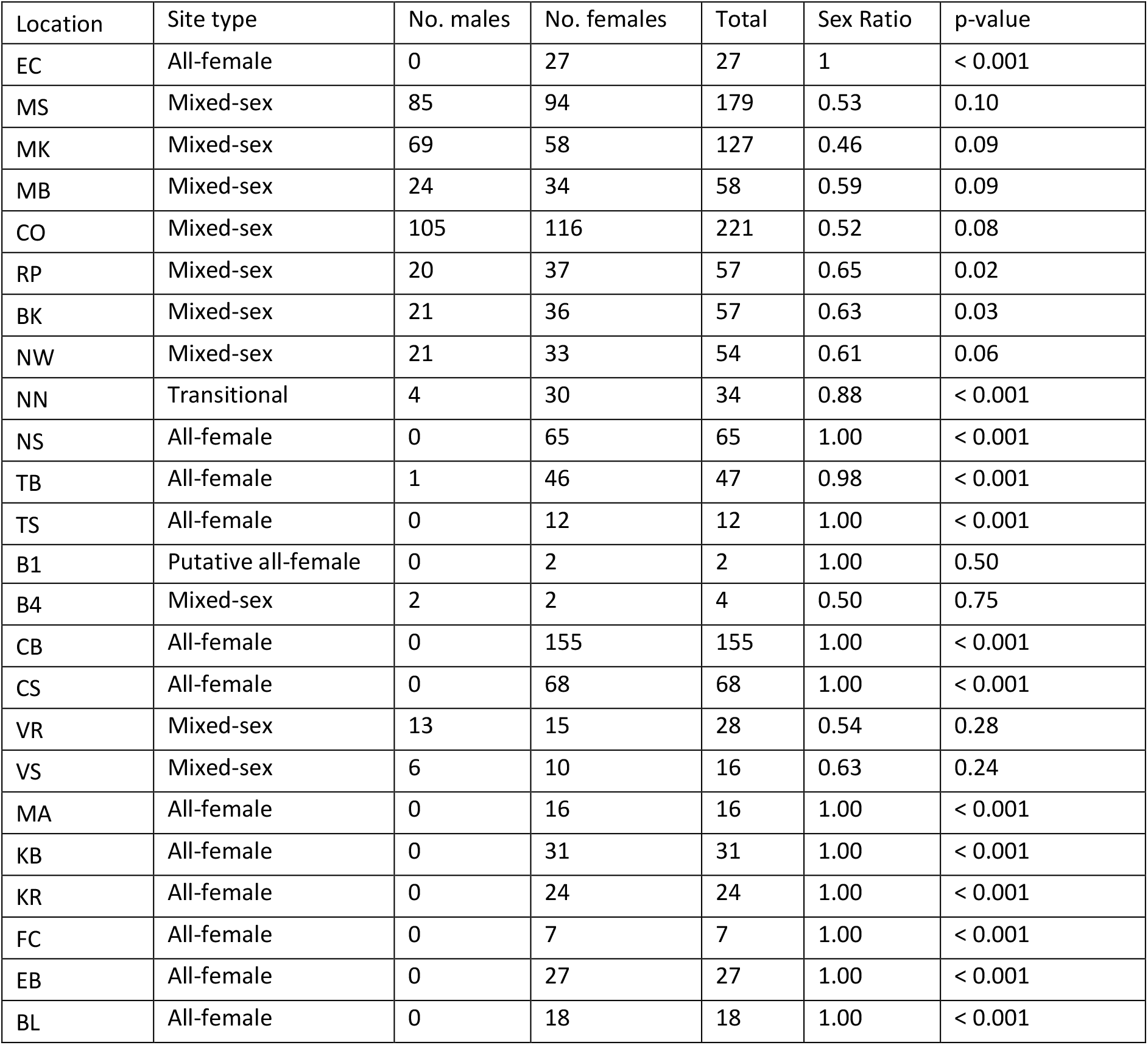
Sample sizes and sex of *M. batesii* adults and nymphs that could be sexed found in the field at each of the surveyed locations (summed over the four years of the study). At sites where we only found females, a binomial two-tailed test was used to determine the probability that it is a mixed-sex location (even sex ratio) based on the number of individuals seen. At sites where males and females have been seen, the location is mixed-sex by definition so we used the binomial test to calculate the probability that the sex ratio of individuals photographed at each location deviated significantly from 50%. At the “transitional” location (NN), only females were seen in 2019-2021 but males were found in 2022.

Eggs collected from most all-female locations produced 100% female hatchlings whereas eggs collected from mixed-sex locations produced approximately equal numbers of female and male hatchlings (see Table S4). We observed only two exceptions to this pattern. A sample of eggs from the all-female site TB produced one hatchling that was phenotypically male-like, and genotyping showed that this hatchling was not a hybrid between locations. This hatchling was therefore either an extremely rare spontaneous male (see Discussion) or a phenotypically atypical female. In addition, at one location (NN) where only females had been observed before 2022, eggs collected from one female in 2021 produced several male hatchlings, and genotyping indicated that this female was either a migrant from the adjacent, genetically similar mixed-sex location BK or that this female had mated with a male migrant from location BK. The following year, adult males were observed at NN and this location therefore appears to be undergoing a transition from all females to mixed-sex.

### Morphology, fecundity, and egg hatching success

We did not find clear and consistent evidence of an effect of reproductive mode across the fitness-related phenotypic traits measured (see Fig. 2b and 2c; for all summary statistics see Table S2). Mean fecundity and egg hatching success were ~12 and 19% lower, respectively, at all-female than at mixed-sex locations, but there was a great deal of variation in these variables among both mixed-sex and all-female locations (Fig. S7). Adult females at all-female locations were more likely to exhibit mild (pMCMC < 0.001) and severe (pMCMC < 0.005) deformities in their non-functional wings than were females at mixed-sex locations (see Fig. S6). Relative to females at mixed-sex locations, adult females at all-female locations had more parasites in *Benstonea*-swamp habitats, but not in *Pandanus*-beach habitats (poptype:habitat interaction: pMCMC < 0.005; see Fig. 3). Post-hoc Tukey tests show this interaction is driven by a near-significant difference between parasitism rates between all-female and mixed-sex sites within swamp habitats (p = 0.054) whereas no other pairwise comparisons approached statistical significance (all p > 0.2; see Table S8).

**Figure 2:**
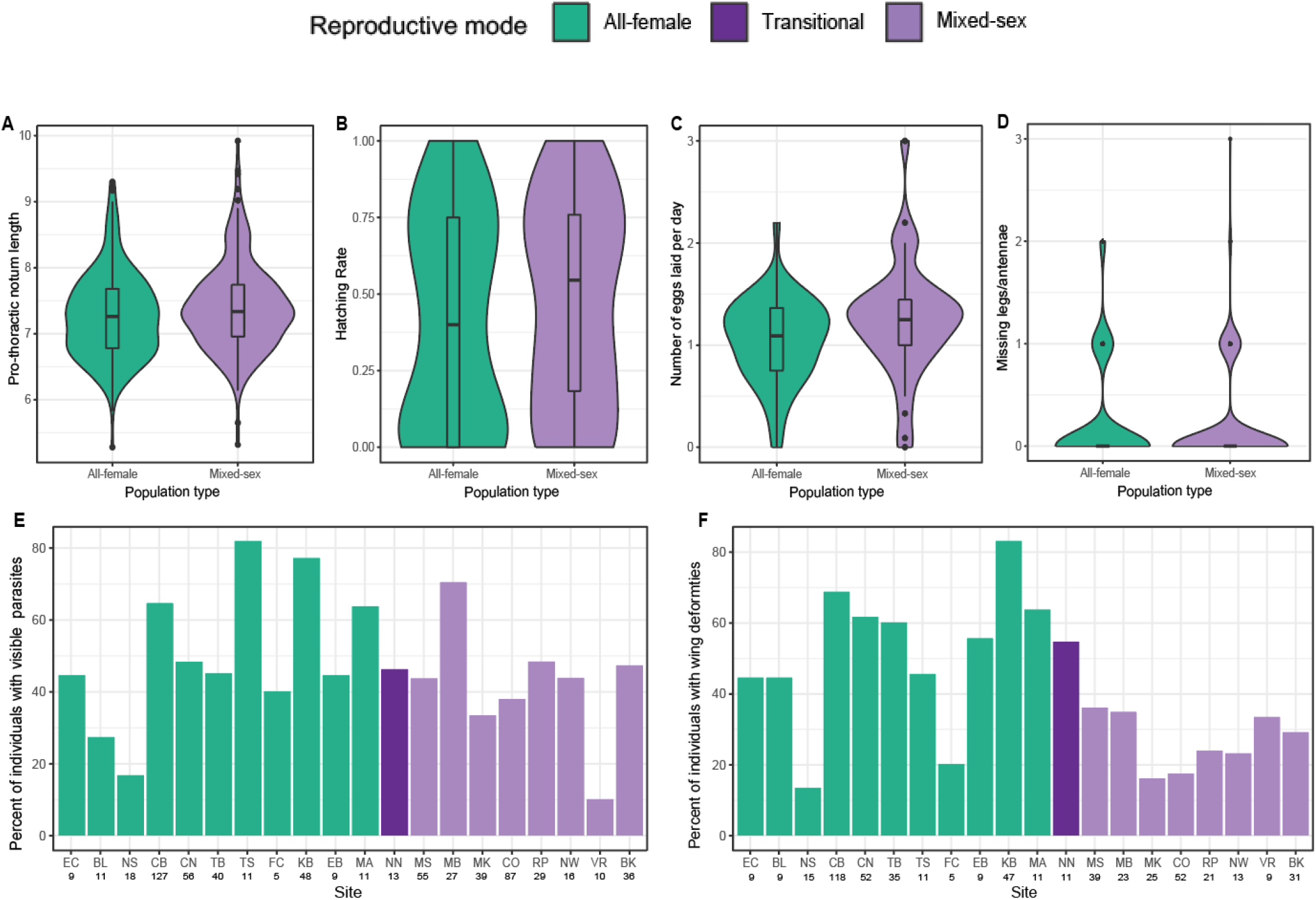
Phenotypic values for *M. batesii* females at mixed-sex and all-female locations: A) Pronotum length as a measure of body size; B) Hatching rate of eggs; C) Number of eggs laid per day as a measure of fecundity; D) Number of missing legs and antennae per individual; E) Percent of adult females observed at each location with visible parasites (fungus, biting midges, or mites); F) Percent of adult females from each location with wing deformities (mild or severe). In panels A – D, means (bars), inter-quartile ranges (boxes) and non-outliner ranges (whiskers) are shown. Violins indicate the distribution of individual values.

**Figure 3:**
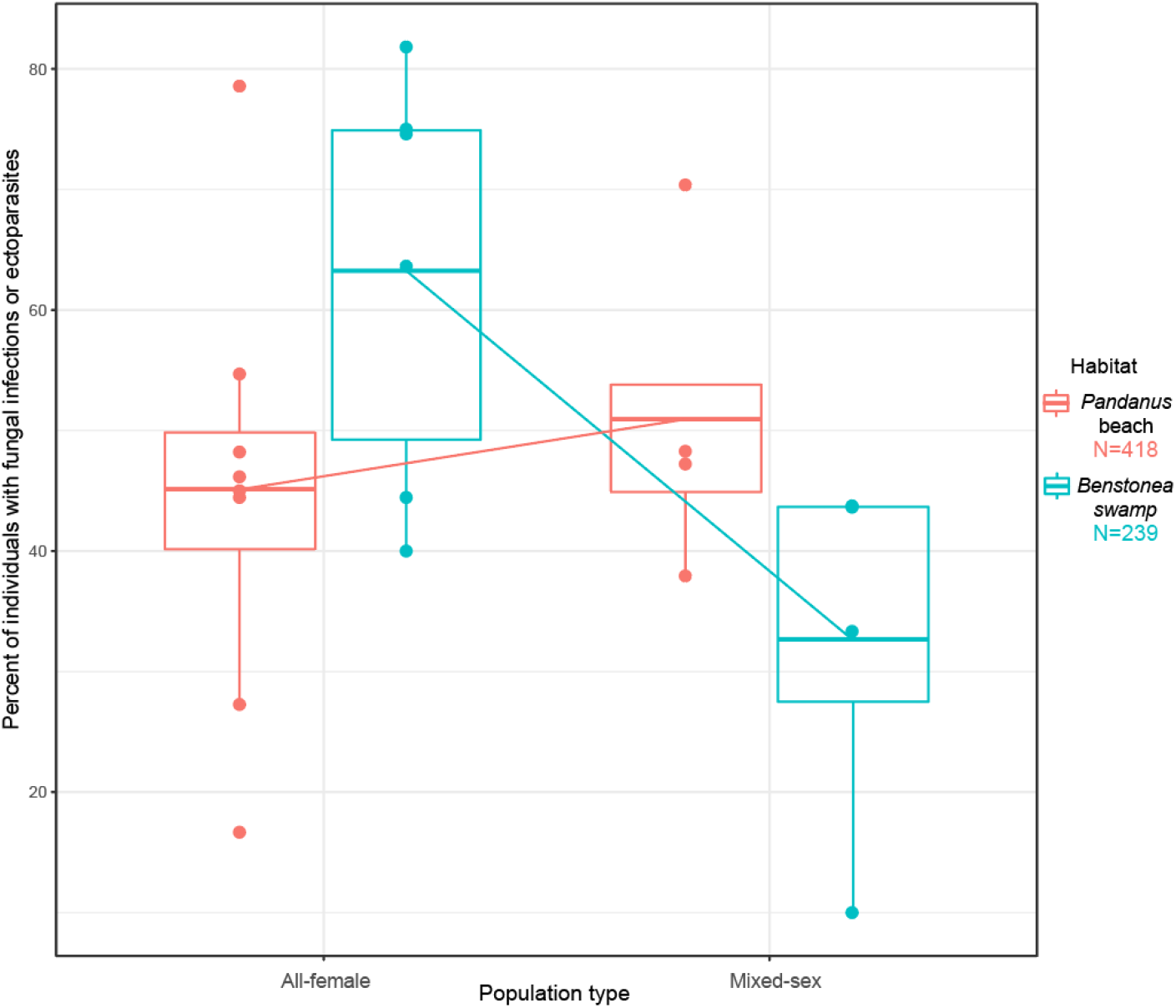
Effects of habitat type and reproductive mode on the rate of visible parasites (fungus, mites or biting midges) in *M. batesii* females at all-female and mixed-sex locations. Each point represents a location mean. Bars indicate means across all populations in each group, boxes represent inter-quartile ranges and whiskers show non-outliner ranges.

**Figure 4:**
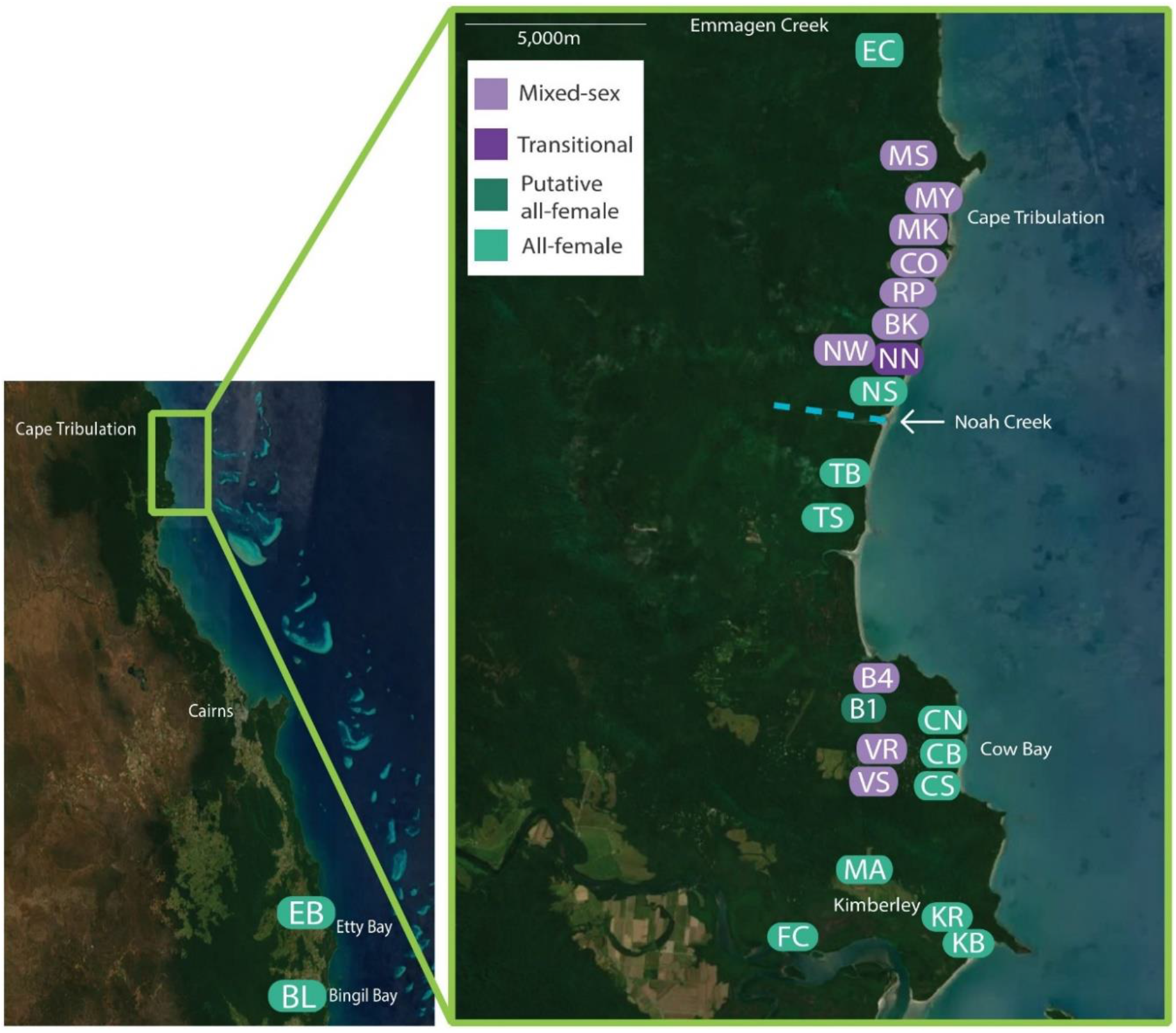
Geographical variation in sex ratio and reproductive mode in *M. batesii*. Two isolated all-female sites occur at the southern limit of the known range of *M. batesii* in Australia, and a spatial mosaic of all-female and mixed-sex sites occurs ~160-180km north of this area between Forest Creek and Emmagen Creek. The blue dashed line represents the mouth of the Noah Creek, which is the main gene flow barrier separating the northern and southern genetic clusters. Locations are characterised as “mixed-sex”, “putative all-female” or “all-female” based on the sex of adults and nymphs photographed at each location (See Table 2). The “transitional” location was all-female in 2019-2021, but males were found at this location in 2022.

Overall, mean rates of parasitism and wing deformities were ~19% and 91% higher, respectively, in females from all-female locations. Additionally, mean numbers of missing legs and antennae were ~40% higher in adult females at all-female locations, but there was no support for an overall difference in this variable between mixed-sex and all-female locations (pMCMC = 0.34). Estimated phylogenetic heritability (H^2^) values suggested effects of genotype for fecundity (H^2^ = 0.74), the presence of deformities (H^2^ = 0.64), the presence of parasites (H^2^ = 0.64), and the hatching success of eggs collected from females at several locations (H^2^ = 0.87). There was moderate support for an effect of genotype on missing antennae and legs (H^2^ = 0.41) but little support for effects of genotype for mild (H^2^ = 0) or severe (H^2^ = 0) wing deformities, or for pronotum length (H^2^ = 0.12).

### Population structure

Sampling sites of *M. batesii* clustered by geographical location rather than by sex ratio. Two major genetic clusters were observed, a “northern” cluster north of the Noah Creek mouth and a “southern” cluster south of the Noah Creek mouth. The Noah Creeks mouth (~ 150 m wide at its widest point) therefore appears to serve as a major barrier to *M. batesii* dispersal and gene-flow. The northern cluster consists mostly of mixed-sex sites while the southern cluster consists mostly of all-female sites, but both genetic clusters contain both all-female and mixed-sex sites (see Fig. 5). Pairwise *F_st_* values reflect the same genetic structure as the cluster analysis, with sites separated by greater geographic distances showing less relatedness (see Table S3). Isolation by distance analysis indicated that genetic differentiation is affected by geographic proximity (Mantel statistic based on Pearson’s product-moment correlation, mantel statistic r = 0.364, p < 0.005). *F_st_* values between the sites varied between 0.16 and 0.96 with lower values between mixed-sex sites in the northern cluster and higher values between all-female sites in the southern cluster.

**Figure 5:**
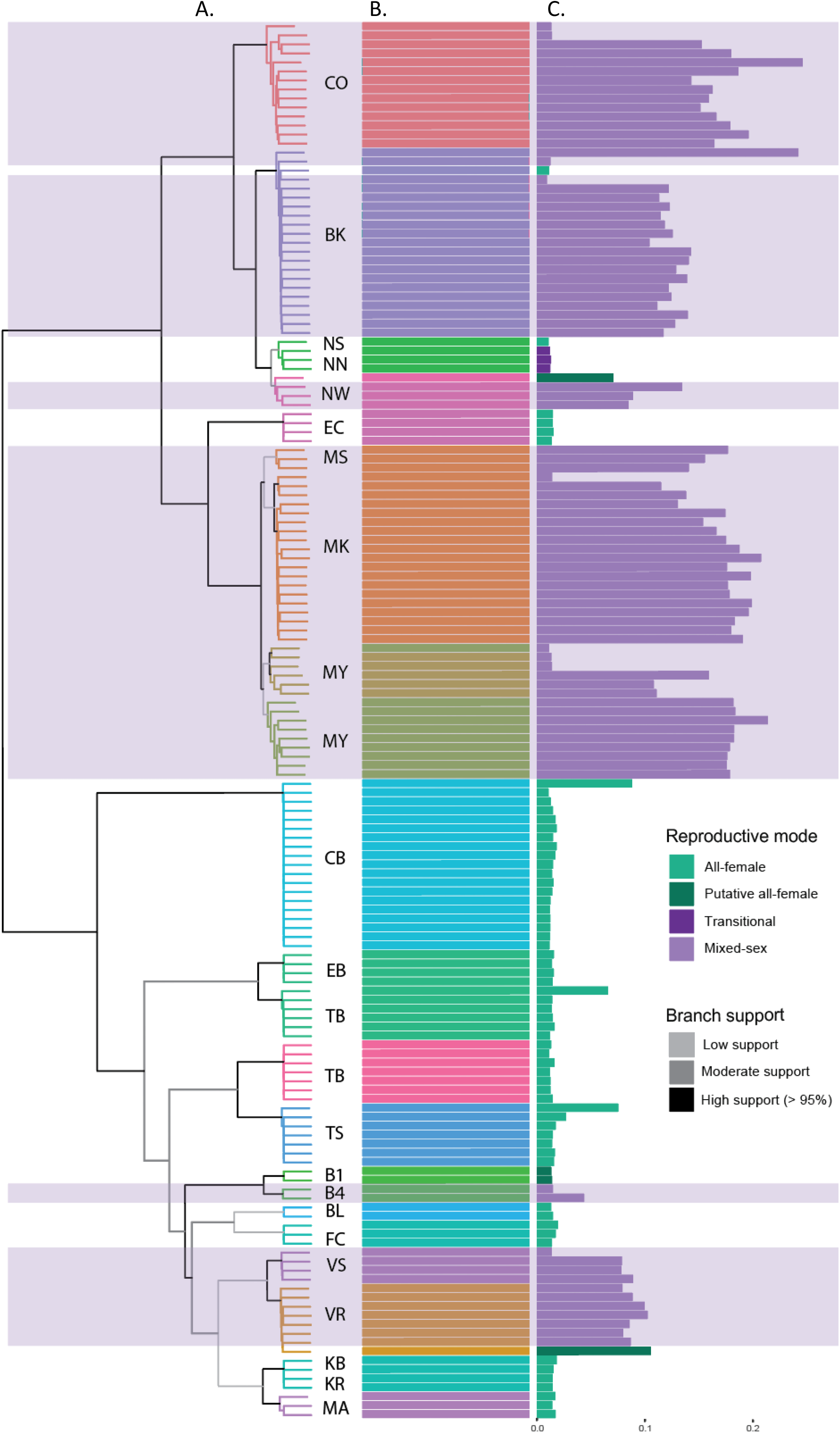
Population structure and heterozygosity of *M. batesii* sites based on 7374 high-quality SNPs from 154 individuals sampled from the field. A) Dendrogram showing relatedness of sampled individuals, with branches colored according to the level of bootstrapped support (based on 10,000 bootstrap iterations) and tips colored to match their highest membership probability. B) Admixture plot showing percent membership in the 17 genetic populations supported by the analysis (Fig. S3); C) Observed heterozygosity values. In plots A and B, different colors represent the primary assigned cluster as indicated by admixture analysis. In plot C, colors represent the observed sex ratio and inferred reproductive mode for each location. Areas shaded in purple show mixed-sex sites.

Admixture analysis revealed that the 155 individuals sampled across the study area belong to 17 genetic populations, with little gene flow detected between sites. Most individuals had 100% membership probability in a single population (Fig. 5b). However, one all-female location (TB) contained multiple genotypes within the one location. Two sets of locations found near one another (< 1000 m) had the same genotype but differing reproductive modes and heterozygosity levels (B1-B4 and NN-NS-NW).

NMDS analysis (Fig. 6) produced a pattern consistent with the population structure obtained from genetic cluster analysis, with the biggest separation found between sites north vs. south of the Noah Creek mouth.

**Figure 6:**
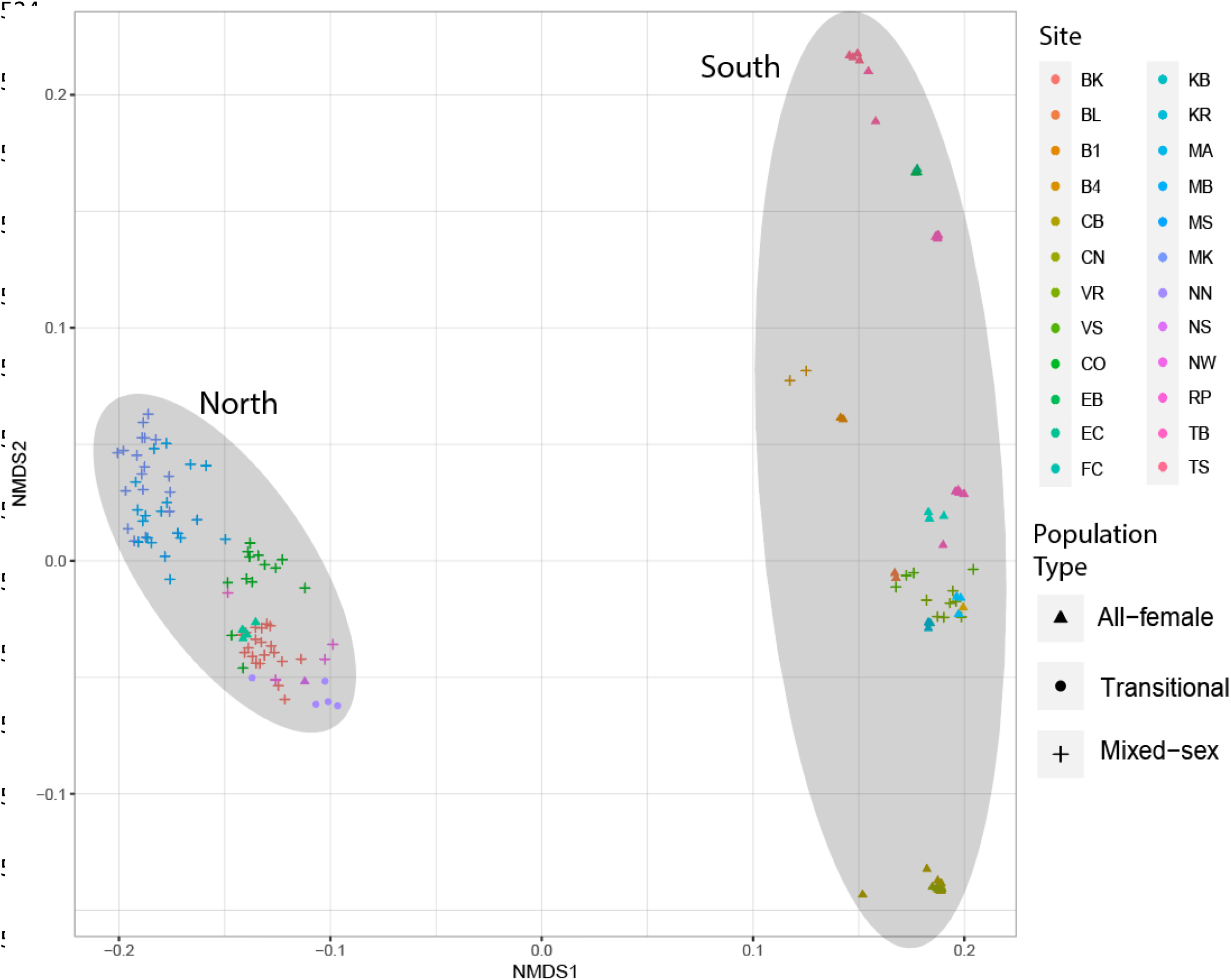
Nonmetric Multidimensional scaling (NMDS) plot of 155 individual *M. batesii* samples based on 7374 high-quality SNPs. Shaded ovals represent individuals found in locations relative to the Noah Creek gene flow barrier.

### Genetic diversity and heterozygosity

Relative to mixed-sex locations, all-female sites had less allelic diversity and Shannon’s diversity, and individuals from all-female sites typically had much lower heterozygosity. All three diversity metrics showed similar patterns across locations (See Fig. S7). There was also substantial variation in genetic diversity between mixed-sex locations: for example, allelic richness was more than 50% higher at CO than at B4. A few individuals sampled from mixed-sex sites had heterozygosity levels comparable to individuals from all-female sites, suggesting that these individuals were produced asexually (see Fig. 5c). On average, heterozygosity of individuals from mixed-sex (sexual) sites was approximately seven times greater than heterozygosity of individuals from all-female (asexual) sites (p < 0.0001: Welsh’s two sample t-test).

## Discussion

We discovered a novel geographical mosaic of all-female and mixed-sex populations of the facultative parthenogen *M. batesii* across its known natural range in Australia. Using this natural system, we carried out the first detailed comparison of both genetic and phenotypic parameters between wild all-female and mixed-sex populations in a facultative parthenogen. Genetic relatedness and admixture indicate multiple independent transitions from sexual reproduction to parthenogenesis. At one location, we also observed a recent re-invasion of males into a previously all-female site. The establishment of female-only locations led to the loss of allelic diversity and a 7-fold reduction in observed heterozygosity, but the phenotypic and fitness consequences of these genetic changes appear to be highly variable among sites. All-female sites had higher rates of mild and severe deformities in their non-functional wings, but did not exhibit increased rates of appendage loss. These sites also had higher rates of fungal and arthropod parasites, but only in *Benstonea*-swamp habitats where parasites may be generally more abundant. However, we found little statistical support for an overall difference in mean body size, fecundity or egg hatching success between all-female and mixed-sex sites. All of the phenotypic traits that we examined varied substantially among both all-female and mixed-sex sites, suggesting that genotype might be more important than mode of reproduction in determining these traits in *M. batesii*, and that high-fitness genotypes are able to thrive without sexual reproduction.

Cluster analyses showed that *M. batesii* populations grouped together based on geographic location, regardless of reproductive mode, and that asexual sites did not form a single monophyletic clade. With one exception (the transition site NN), this pattern remained stable over four years of sampling. We can infer at least four independent transitions from sexual to parthenogenetic reproduction, resulting in all-female sites: the assumed transition from a sexually reproducing ancestral population to an all-female population that gave rise to the genetic cluster of asexual populations south of the gene-flow barrier at Noah Creek, plus three more recent transitions resulting in all-female populations EC, NS, and B1, all of which are very close spatially and genetically to adjacent mixed-sex sites. Admixture analysis showed little gene flow between the locations despite their geographical proximity (see Fig. 5b). It is not known how long ago these reproductive mode transitions occurred, because no historic samples or previous literature is available that could be used to quantify time of establishment. However, the close geographical proximity and genetic relatedness of some sexual and asexual sites suggest recent establishment. For example, sites B1 and B4 as well as NW and NS exhibit differing reproductive modes, but they are still considered to be from the same population based on admixture analysis. Additionally, results of inter-population crosses (to be presented in a separate paper) show that individuals from sexual and asexual sites are interfertile and can therefore be regarded as a single species.

We identified two main genetic groups, separated by a large estuary (Noah Creek) that serves as a gene flow barrier. The northern group consists of mainly sexual sites and the southern group consists of mainly asexual sites. This partially resembles the geographic pattern found in other geographically parthenogenetic phasmids in the southern hemisphere (Buckley, Marske & Attanayake, 2009; Morgan-Richards, Trewick & Stringer, 2010), and contrasts with the pattern in the northern hemisphere (Law & Crespi, 2002). However, the northernmost site that we sampled in this study (EC) is all-female, while some southern populations (VS, VR, B4) are mixed-sex. Thus, *M. batesii* appears to exhibit a mosaic of sexual and asexual populations across its known range in Australia, rather than a clear latitudinal gradient. Asexually reproducing populations of species that display geographical parthenogenesis are often found in marginal habitats, typically at a higher altitude or latitude than sexually reproducing populations (Lynch, 1984; Kearney, 2005; Tilquin & Kokko, 2016). However, the locations examined in our study are found in similar tropical coastal-lowland rainforest habitats spanning a small latitudinal range (0.24°; 1.8° including the isolated southern populations), with little variation in temperature, humidity, or precipitation between locations. Additionally, sexual and asexual sites were both found in each of the two major habitat types (*Pandanus*-beach and *Benstonea*-swamp), so climate or habitat differences do not appear to explain the variation in reproductive mode.

The lack of obvious environmental barriers or differences in habitat between the sexual and asexual sites suggests that this spatial pattern of variation in reproductive mode results from factors intrinsic to the biology of *M. batesii*. Four intrinsic processes might have contributed to the observed spatial pattern: dispersal and colonization of new habitats by unmated females, male invasion of established all-female populations, spontaneous male genesis from unfertilized eggs, and local losses of sex through male extinction.

First, if females disperse more than males, they could travel further from the mixed-sex sites and establish all-female sites that could avoid male invasion for many generations. All-female sites could persist once established if parthenogenetically produced females exhibit reduced propensity of females to mate, and reduced benefits of mating, as reported for the related phasmid *Extatosoma tiaratum* (Burke & Bonduriansky, 2022). However, this explanation seems implausible because *M. batesii* adults of both sexes of this species disperse similar distances (Boldbaatar, 2022).

Second, if sexual sites south of the gene-flow barrier (VR, VS, B4) arose as a result of males from the north invading southern all-female sites, then these sites would show a genetic admixture pattern consistent with hybrid origin. We found no such pattern. Rather, all the southern sites clustered together, and none showed evidence of genetic admixture from northern sites. However, we observed an apparent on-going transition from all-female to mixed-sex at a northern site (NN), showing that male invasion of all-female sites is possible and might be relatively frequent at locations where all-female sites arise in close proximity to mixed-sex sites.

Third, sex may re-invade after it is lost through the spontaneous production of males from unfertilized eggs. The sexual sites in the south (VS, VR, B4) could have arisen via such spontaneous male genesis. Spontaneous male genesis is possible in diploid species with XX XO sexual karyotypes; due to a meiotic/developmental error, the second X chromosome can be lost in situ in parthenogenically produced offspring, resulting in a genetic male (Pijnacker & Ferwerda, 1980; Scali, 2009). The viability and reproductive success of spontaneous males has not been studied in any phasmid species, so the ability for spontaneous males to successfully turn an established asexual population to sexual is currently unknown (see Morgan-Richards, Langton-Myers & Trewick, 2019). However, in the obligate asexual marine snail *Potamopyrgus antipodarum*, low rates of male production and reduced functionality in asexually-produced males were found to make male invasion in asexual populations unlikely (Neiman et. al., 2012; Jalinsky, Logsdon Jr., & Neiman, 2020) The sexual karyotype of *M. batesii* is currently unknown, but diploid phasmids are known to have either XX XY or XX XO sexual karyotypes (Scali, 2009; Schwander & Crespi, 2009), making spontaneous male genesis a possibility. If spontaneous males are possible in *M. batesii*, they are extremely rare (< 1/1000 parthenogenetic eggs). Out of 533 eggs from all-female sites that were collected in the field, we found one morphologically male-like hatchling, although morphologically ambiguous hatchlings are occasionally found. This hatchling was not reared to adulthood, so it was not conclusively determined to be male. Out of 1083 of eggs produced parthenogenetically in the lab, no males were found. Thus, the spontaneous re-emergence of sex in female-only locations appears unlikely in this species.

Fourth, female-only sites could result from local extinction of males. This seems the most likely explanation for the geographic mosaic of sexual and asexual localities that we report: northern all-female sites (EC, NS) could have resulted from male extinction, while southern mixed-sex sites (VS, VR, B4) could represent remnants of an ancestral mixed-sex population south of Noah Creek. Male extinction could occur stochastically in *M. batesii* populations, and such a pattern of extinction may be especially likely in small, isolated populations (Pianka, 2000). Male extinction might also occur as a result of the evolution of female resistance to mating. There are many costs of sexual reproduction (Lehtonen, Jennions & Kokko, 2012; but see Otto & Lenormand, 2002). In a facultatively parthenogenetic species, such costs could generate sexual conflict and favour female resistance to mating. Sexual conflict can play a large role in sexual/asexual population dynamics, involving evolution of male traits that harm females and female counter-adaptations that minimize the harm (Kawatsu, 2013; Gerber & Koko, 2016; Burke & Bonduriansky, 2019). *Megacrania batesii* males guard females in mixed-sex sites for two weeks or more, and most adult females at these sites carry guarding males (Boldbaatar, 2022). This guarding behavior could interfere with female foraging, reducing fitness of *M. batesii* females. Gerber and Kokko (2016) found that female resistance tactics are most beneficial at low population densities, which occurs in many of the sites described in this paper. Thus, opportunity for the evolution of female resistance tactics appears to exist in this species. However, modeling by Kawatsu (2013) found that antagonistic evolution driven by sexual conflict ultimately leads to the prevalence of sex or the co-occurrence of both reproductive modes unless strong reproductive isolation evolves. *Megacrania batesii* investigated in this study do not show complete reproductive isolation across sites: females from asexual locations can mate with males from mixed-sex locations and produce sons when paired in the lab (to be presented in a separate paper). However, we found that some individuals from sexual sites have heterozygosity values comparable to those of asexual sites. This suggests that females from mixed-sex sites occasionally reproduce asexually despite an abundance of males. It is possible that some females fail to mate or experience fertilization failure after mating, and selection favouring asexual reproduction could potentially lead to the evolution of female mating avoidance or resistance to fertilisation (Burke & Bonduriansky, 2022).

We found a striking contrast in genetic diversity between the sexual and asexual populations. Asexual sites had on average 85% less allelic richness, 85% less Shannon’s diversity, and 83.5% less heterozygosity than their sexual counterparts. This dramatic difference in genetic diversity could be explained by the mechanism of parthenogenesis in *M. batesii*. The cytological mechanisms underlying asexual reproduction determines how much heterozygosity is maintained when restoring the parental chromosome number (Stenberg & Saura, 2009; Moritz et al. 1990). Preliminary results suggest that parthenogenesis in *M. batesii* results in substantial (but not complete) loss of heterozygosity over a single generation of parthenogenetic reproduction, consistent with automixis via either terminal or central fusion (Stenberg & Saura, 2009).

Yet, we found no consistent overall effect of reproductive mode on body size, number of missing appendages, or female reproductive output. Average fecundity and egg viability were 12% and 19% lower, respectively, in all-female sites compared to mixed-sex sites, but there was no statistical support for an effect of reproductive mode on these traits given the considerable variation between sites within each reproductive mode. These findings suggest that, while asexual reproduction could negatively affect some aspects of performance, other factors such as inbreeding in smaller sexual populations or the prevalence of highly locally adapted genotypes in some asexual populations could affect performance to a comparable degree.

*M. batesii* from asexual sites had higher rates of mild and severe wing deformities, in addition to large inter-site variation in these traits (see Fig. 2d and 2e). Low heterozygosity could explain the higher rates of wing deformities in all-female sites. Parthenogenetically produced *Extatosoma tiaratum* females exhibit higher rates of wing deformities than do sexually produced females (Burke & Bonduriansky 2022). Likewise, multiple homozygous loss of function mutations cause wing defects in *Drosophila* (Terriente-Felix, López-Varea & de Celis, 2010; George et. al., 2019). If *M. batesii* shares any of these genes, asexually produced females may be more at risk of displaying a homozygous genotype resulting in the loss of function of certain genes relating to wing formation during development. Additionally, the high rates of wing deformities in asexual sites may reflect developmental instability as seen with wing asymmetry in pea aphids (Hammelman et. al., 2020). While wing deformities may be more prevalent in asexual sites, both sexes of *M. batesii* are flightless and wing deformities might have little or no effect on fitness.

We also found elevated rates of fungal and arthropod parasitism of females in all-female sites when compared to mixed-sex sites within *Benstonea*-swamp habitats but not within *Pandanus*-beach habitats. Many *M. batesii* individuals had brown or black spots on their head, thorax, abdomen, or legs that appeared to be a fungal infection, and some individuals carried mites or biting midges. Elevated rates of parasites in asexual lineages are consistent with Red Queen dynamics, such that sexual populations are better equipped than asexual populations in evolutionary arms races against parasites due to their higher levels of recombination and genotypic diversity (Hamilton, Axelrod & Tanese, 1990; Hamilton & Zuk, 1982). Indeed, Hite (2017) and Lively (1987) found higher frequencies of males and sexually produced offspring in the sea snail, *Dentistyla dentifera*, and the New Zealand mud snail, *Potomopyrgus antipodarum* respectively, during fungal epidemics. Some asexual *M. batesii* sites (KB, CB, and TS) may be more susceptible to parasites due to their low heterozygosity levels but these patterns were not consistent across all asexual sites. Instead, we found an interaction between reproductive mode and habitat type, suggesting that asexual reproduction leads to elevated rates of infections for *M. batesii* found in closed-canopy rainforest swamps on *Benstonea* host plants, but not in *Pandanus*-dominated beach habitat. This might be because high humidity in swamp habitats allows for more fungal abundance and growth. Our results therefore suggest that asexual *M. batesii* populations are disadvantaged in their ability to cope with parasites only in certain parts of their range.

Our finding of considerable among-site variation in performance suggests that some asexual *M. batesii* lineages have genotypes that are well-adapted to their environment and thus perform well despite extremely low heterozygosity and allelic diversity. Thus, the loss of sex and consequent genetic changes may not be sufficient to drive asexual lineages to extinction. Rather, locally adapted asexual lineages might persist and even thrive, perhaps for many generations. Ultimately, however, it is possible that outside factors, like changes in climate or the invasion of new competitors or parasites, might impose strong selection, and asexual populations might suffer extinction as a result of failure to adapt to such changes.

Understanding the consequences of sexual vs. asexual reproduction in *M. batesii* is also relevant to conservation because this species has a restricted natural range and could be quite vulnerable to global heating, habitat loss, or novel pathogens. Cermak and Hasenpusch (2000) designated this species as vulnerable due to its range being less than 100km^2^, but also noted that if the populations were found to be genetically distinct they may represent declining remnants of larger ancestral populations, and therefore conservation management should be considered. Indeed, we did find that the genetic groups were spatially isolated from one another. This puts them more at risk of extinction events as small, isolated populations are more likely to undergo Muller’s ratchet and accumulate deleterious mutations (Muller, 1964). Asexual populations might be especially at risk of extinction in changing environments. The importance of field-based insect studies increases as concerns grow about a substantial, ongoing decline in terrestrial insect abundances around the world (van Klink et. al., 2020). Long-term monitoring of natural insect populations could help to better estimate the level of threat to their survival and to inform future management.

## Notes

### Competing Interest Statement

The authors have declared no competing interest.

### Summary of Updates

The title page has been stripped of the author names and affiliations. A post-hoc Tukey test was added.

