## Supplemental Tables and Figures for "Genetic and phenotypic consequences of local transitions between sexual and parthenogenetic reproduction in the wild"

### Supplementary Tables and Figures:

Table S1. Results of MCMCglmm modelling of phenotypic variables from individuals in natural populations of *M. batesii* (see Table 1 in main text for parameters). Green highlight indicates p-values lower than 0.05. Mode of reproduction differentiates mixed-sex populations from all-female populations. Habitat type differentiates *Pandanus*-beach from *Benstonea*-swamp habitats. All models included a matrix of genetic relatedness and year of observation as random effects.

| | Posterior Mean | l-95% CI | u-95% CI | Effective sample size | $H^2$ Phylogenetic Heritability | l-95% CI $H^2$ | u-95% CI $H^2$ | pMCMC |
|---|---|---|---|---|---|---|---|---|
| **Pronotum Length** | | | | | 0.1239 | 0.05136 | 0.4067 | |
| Mode of reproduction | 0.05138 | -0.4601 | 0.474 | 1990 | | | | 0.771 |
| Habitat type | 0.02439 | -0.162 | 0.216 | 1990 | | | | 0.786 |
| Mode x Habitat | 0.11536 | -0.329 | 0.535 | 1990 | | | | 0.622 |
| **Fungus/Parasites** | | | | | 0.639 | 0.0420 | 0.3909 | |
| Mode of reproduction | -0.5075 | -1.616 | 0.5637 | 1980 | | | | 0.341 |
| Habitat type | -0.5894 | -1.545 | 0.450 | 1980 | | | | 0.2131 |
| Mode x Habitat | 1.571 | 0.4877 | 2.737 | 1980 | | | | 0.004 |
| **Missing Antennae/Legs** | | | | | 0.41 | 0.186 | 0.73 | |
| Mode of reproduction | 0.325 | -0.41 | 0.97 | 1654 | | | | 0.339 |
| Habitat type | -0.041 | -0.499 | 0.381 | 1654 | | | | 0.857 |
| **Mild wing deformities** | | | | | -6.574e-06 | 8.439e-07 | 0.0036 | |
| Mode of reproduction | 128.85 | 31.44 | 237.31 | 2946 | | | | 0.0006 |
| Habitat type | 25.51 | -76.61 | 135.50 | 3507 | | | | 0.611 |
| Mode x Habitat | -18.67 | -156.43 | 115.94 | 3314 | | | | 0.773 |
| **Severe wing Deformities** | | | | | 3.445e-06 | 5.264e-07 | 2.288e-05 | |
| Mode of reproduction | 167.66 | 36.88 | 299.39 | 2673 | | | | 0.00483 |
| Habitat type | 36.05 | -80.15 | 148.46 | 3314 | | | | 0.47314 |
| Mode x Habitat | 15.49 | -109.60 | 154.01 | 3314 | | | | 0.80628 |
| **Fecundity (eggs per day)** | | | | | 0.737 | 0.389 | 0.940 | |
| Mode of reproduction | -0.015 | -1.153 | 1.237 | 2990 | | | | 0.978 |
| **Egg Hatching Success** | | | | | 0.872 | 0.525 | 0.959 | |
| Mode of reproduction | 0.724 | -1.987 | 3.535 | 3300 | | | | 0.601 |
| Habitat type | 0.449 | -2.161 | 2.894 | 3300 | | | | 0.713 |
| Mode x Habitat | 0.226 | -2.517 | 2.929 | 3300 | | | | 0.880 |

Table S2: Summary statistics and sample sizes for the traits included in the phenotypic analyses. Phenotypic values were determined from photos of *M. batesii* individuals on their host-plants at multiple locations across the study area. Habitat is categorised as *Pandanus* sp. host-plants along the beach margin or *Benstonea* sp. host-plants in closed-canopy rainforest swamp.

| | | | Pronotum | | | Fungus/ Parasites | | | Fecundity Eggs/day | | | Hatching Rate | | |
|---|---|---|---|---|---|---|---|---|---|---|---|---|---|---|
| **Site** | **Type** | **Habitat** | n | mean | sd | n | presence | rate | n | mean | sterr | n | mean | std err |
| BK | Mixed-sex | beach | 33 | 7.41 | 0.74 | 36 | 17 | 0.47 | 18 | 1.22 | 0.27 | 205 | 0.44 | 0.03 |
| BL | All-female | beach | 7 | 6.84 | 0.65 | 11 | 3 | 0.27 | | | | | | |
| B1 | Putative All-female | swamp | 2 | 7.17 | 0.16 | 2 | 0 | 0 | | | | | | |
| B4 | Mixed-sex | swamp | 2 | 7.72 | 1.62 | 2 | 2 | 1 | | | | | | |
| CB | All-female | beach | 25 | 7.62 | 0.65 | 64 | 35 | 0.55 | 22 | 1.03 | 0.37 | 310 | 0.58 | 0.03 |
| CN | All-female | beach | 51 | 7.25 | 0.55 | 56 | 27 | 0.48 | | | | | | |
| CS | All-female | swamp | 58 | 7.56 | 0.61 | 63 | 47 | 0.75 | | | | | | |
| VR | Mixed-sex | swamp | 10 | 6.96 | 0.47 | 10 | 1 | 0.1 | 1 | 0.33 | | 8 | 0.88 | 0.13 |
| VS | Mixed-sex | swamp | 3 | 7.65 | 0.04 | 3 | 2 | 0.67 | | | | | | |
| CO | Mixed-sex | beach | 83 | 7.4 | 0.64 | 87 | 33 | 0.38 | 5 | 1.10 | 0.41 | 61 | 0.70 | 0.06 |
| EB | All-female | beach | 7 | 7.53 | 0.59 | 9 | 4 | 0.44 | | | | | | |
| EC | All-female | swamp | 3 | 7.44 | 0.21 | 9 | 4 | 0.44 | | | | | | |
| FC | All-female | swamp | 2 | 7.69 | 0.45 | 5 | 2 | 0.4 | | | | | | |
| KB | All-female | beach | 25 | 6.87 | 0.58 | 28 | 22 | 0.79 | 4 | 1.22 | 0.28 | 30 | 0.1 | 0.06 |
| KR | All-female | swamp | 21 | 6.74 | 0.54 | 20 | 15 | 0.75 | | | | | | |
| MA | All-female | swamp | 6 | 7.68 | 0.59 | 11 | 7 | 0.64 | | | | | | |
| MB | Mixed-sex | beach | 20 | 7.38 | 0.54 | 27 | 19 | 0.7 | 8 | 1.17 | 0.25 | 152 | 0.57 | 0.04 |
| MS | Mixed-sex | swamp | 15 | 7.83 | 0.87 | 55 | 24 | 0.44 | | | | | | |
| MK | Mixed-sex | swamp | 36 | 7.33 | 0.67 | 39 | 13 | 0.33 | 8 | 1.58 | 1.05 | 31 | 0.74 | 0.08 |
| NN | All-female | beach | 2 | 7.86 | 0.32 | 13 | 6 | 0.46 | 3 | 1.47 | 0.64 | | | |
| NS | All-female | beach | | | | 18 | 3 | 0.17 | | | | | | |
| NW | Mixed-sex | swamp | 15 | 7.67 | 0.63 | 16 | 7 | 0.44 | | | | | | |
| RP | Mixed-sex | beach | 27 | 7.289 | 0.54 | 29 | 14 | 0.48 | | | | | | |
| TB | All-female | beach | 27 | 7.22 | 0.61 | 40 | 18 | 0.45 | 12 | 0.93 | 0.44 | 130 | 0.58 | 0.04 |
| TS | All-female | swamp | 11 | 6.77 | 0.37 | 11 | 9 | 0.82 | | | | 205 | 0.44 | 0.03 |

| | | Mild Wing Deformities | | | Severe Wing Deformities | | | Missing Antennae/Legs | | | | |
|---|---|---|---|---|---|---|---|---|---|---|---|---|
| **Population** | **Type** | n | presence | rate | n | presence | rate | **n** | **presence** | **rate** | **mean** | std err |
| BK | Mixed-sex | 31 | 6 | 0.19 | 31 | 3 | 0.10 | 36 | 5 | 0.14 | 0.14 | 0.06 |
| BL | All-female | 9 | 2 | 0.22 | 9 | 2 | 0.22 | 10 | 1 | 0.10 | 0.20 | 0.20 |
| B1 | Putative All-female | 1 | 0 | 0 | 1 | 0 | 0 | 2 | 1 | 0.50 | 1.00 | 1.00 |
| B4 | Mixed-sex | 2 | 1 | 0.5 | 2 | 1 | 0.5 | 2 | 0 | 0.00 | 0.00 | 0.00 |
| CB | All-female | 78 | 21 | 0.27 | 78 | 30 | 0.38 | 139 | 21 | 0.15 | 0.16 | 0.03 |
| CN | All-female | 57 | 18 | 0.32 | 57 | 16 | 0.28 | 57 | 14 | 0.25 | 0.30 | 0.07 |
| CS | All-female | 62 | 16 | 0.26 | 62 | 30 | 0.48 | 18 | 3 | 0.17 | 0.17 | 0.09 |
| VR | Mixed-sex | 9 | 3 | 0.33 | 9 | 0 | 0 | 10 | 4 | 0.40 | 0.40 | 0.16 |
| VS | Mixed-sex | 2 | 0 | 0 | 2 | 1 | 0.5 | 4 | 0 | 0.00 | 0.00 | 0.00 |
| CO | Mixed-sex | 52 | 7 | 0.13 | 52 | 2 | 0.04 | 95 | 11 | 0.12 | 0.12 | 0.03 |
| EB | All-female | 9 | 1 | 0.11 | 9 | 4 | 0.44 | 9 | 4 | 0.44 | 0.56 | 0.24 |
| EC | All-female | 9 | 3 | 0.33 | 9 | 1 | 0.11 | 10 | 2 | 0.20 | 0.30 | 0.21 |
| FC | All-female | 5 | 1 | 0.2 | 5 | 0 | 0 | 5 | 1 | 0.20 | 0.20 | 0.20 |
| KB | All-female | 28 | 9 | 0.32 | 28 | 13 | 0.46 | 27 | 10 | 0.37 | 0.41 | 0.11 |
| KR | All-female | 22 | 9 | 0.41 | 22 | 10 | 0.45 | 22 | 6 | 0.27 | 0.32 | 0.12 |
| MA | All-female | 11 | 4 | 0.37 | 11 | 3 | 0.27 | 11 | 3 | 0.27 | 0.36 | 0.20 |
| MB | Mixed-sex | 23 | 3 | 0.13 | 23 | 5 | 0.22 | 28 | 7 | 0.25 | 0.29 | 0.10 |
| MS | Mixed-sex | 39 | 9 | 0.23 | 39 | 5 | 0.13 | 60 | 11 | 0.18 | 0.22 | 0.06 |
| MK | Mixed-sex | 25 | 1 | 0.04 | 25 | 3 | 0.12 | 50 | 10 | 0.20 | 0.26 | 0.08 |
| NN | All-female | 11 | 3 | 0.27 | 11 | 3 | 0.27 | 21 | 4 | 0.19 | 0.24 | 0.12 |
| NS | All-female | 15 | 1 | 0.07 | 15 | 1 | 0.07 | 50 | 5 | 0.10 | 0.12 | 0.05 |
| NW | Mixed-sex | 13 | 1 | 0.08 | 13 | 2 | 0.15 | 20 | 4 | 0.20 | 0.20 | 0.09 |
| RP | Mixed-sex | 21 | 2 | 0.10 | 21 | 3 | 0.14 | 27 | 4 | 0.15 | 0.15 | 0.07 |
| TB | All-female | 35 | 13 | 0.37 | 35 | 8 | 0.23 | 42 | 12 | 0.29 | 0.36 | 0.10 |
| TS | All-female | 11 | 2 | 0.18 | 11 | 3 | 0.27 | 12 | 3 | 0.25 | 0.25 | 0.13 |

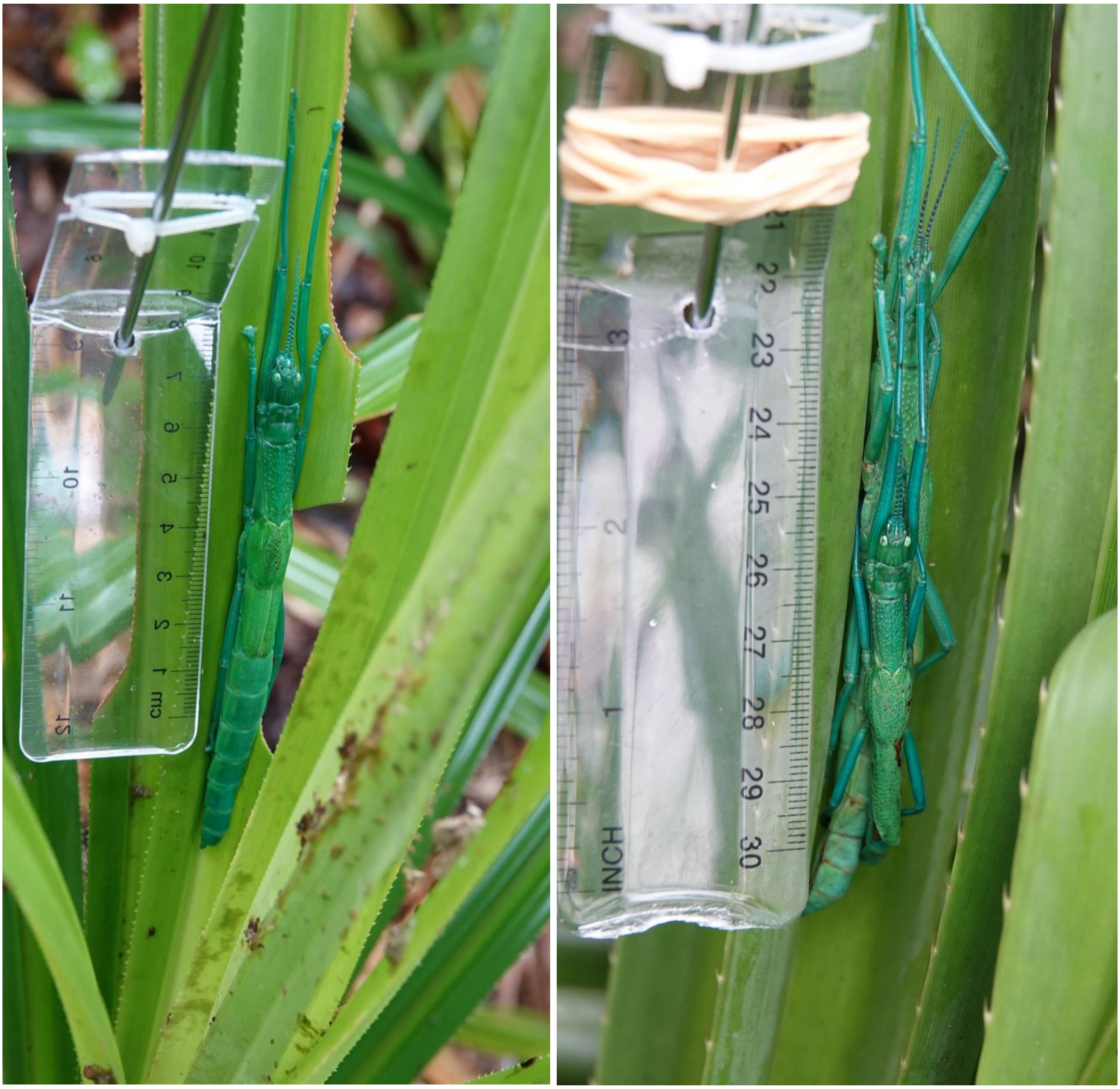


Fig. S1. *Megacrania batesii* individuals and pairs were photographed on their host plants alongside a scale.

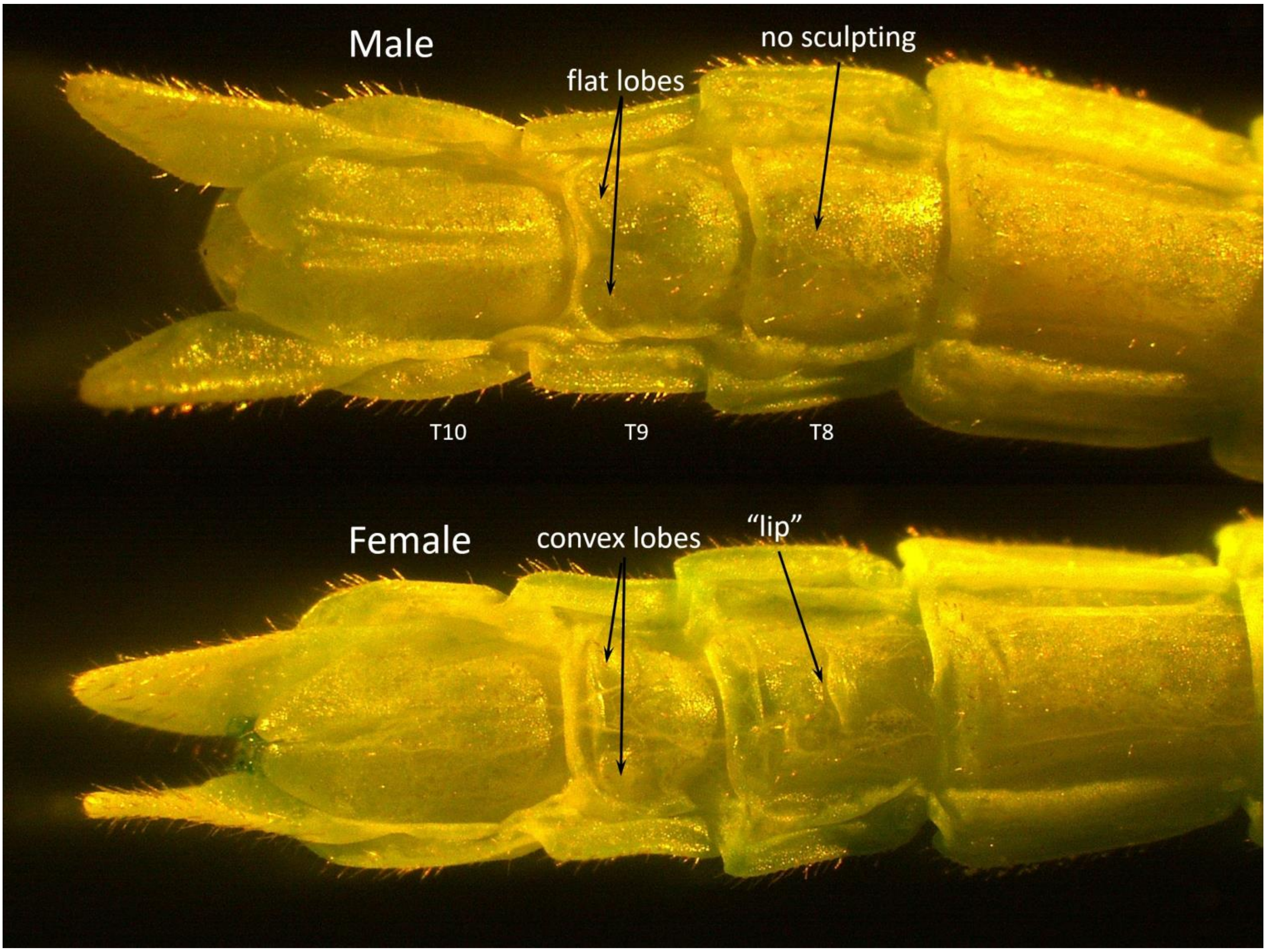


Fig. S2. Ventral view of female (bottom) and male (top) *Megacrania batesii* hatchling abdomen tip (segments T8, T9, T10), showing sexual dimorphism in sternite morphology. Females have a prominent “lip” on the 8th sternite and two convex, pointed lobes on the 9th sternite. Males have a smooth (unsculpted) 8th sternite and two flattened, arch-shaped lobes on the 9th sternite.

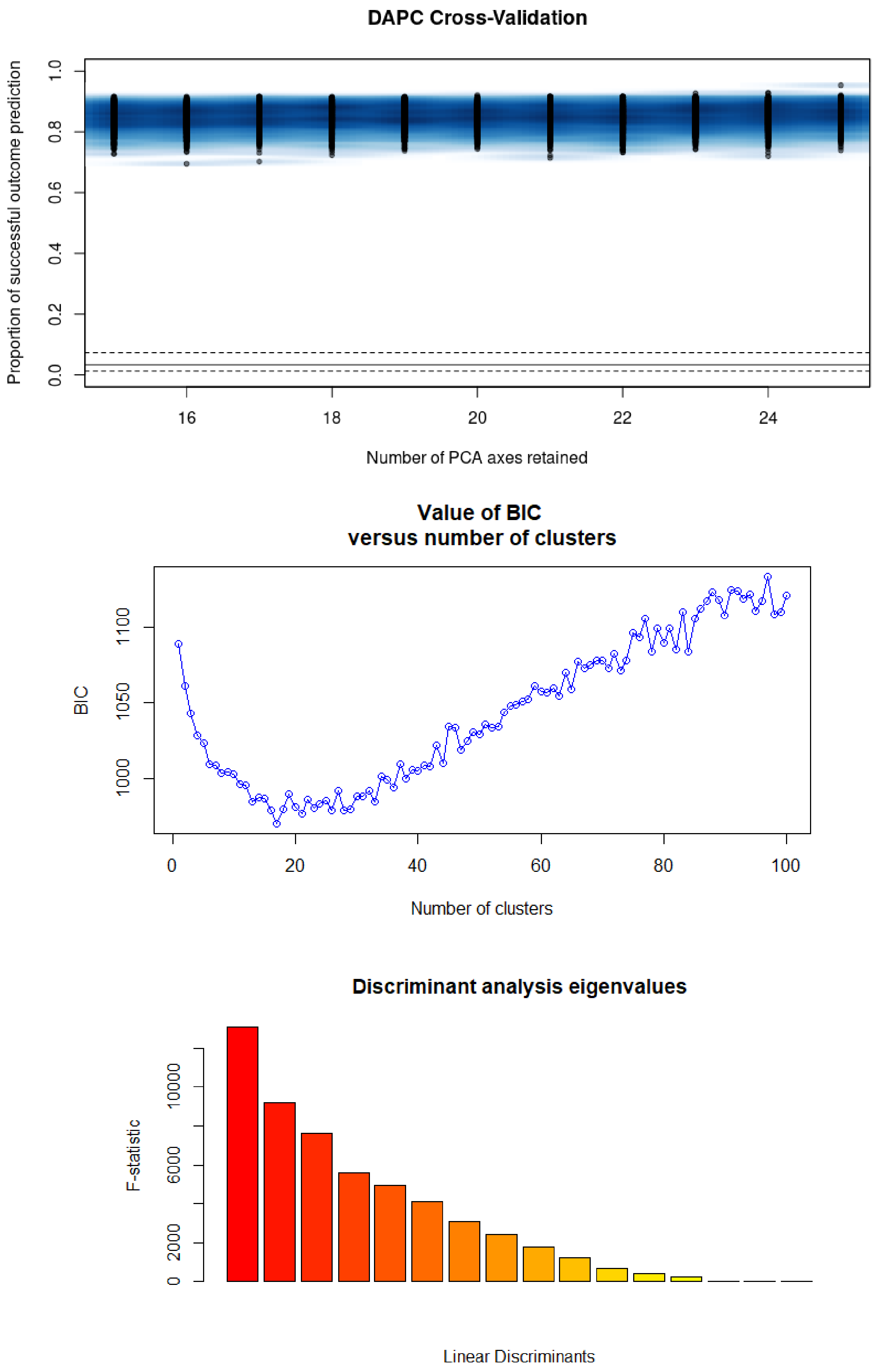


Fig. S3. BIC values and DA eigenvalues. 17 genetic populations were determined based on lowest consistent BIC score and 6 discriminants were chosen based on the number of discriminants that, combined, made up most of the variance.

Table S3: Pairwise Fst matrix with Fst values on lower left cells of the dividing line and p-values in upper right cells. Cells with values indicating non-significant Fst between populations are shaded red.

| | BK | MK | TB | BL | CO | CB | KR | B4 | KB | B1 | NN | VR | EB | MB | MS | EB | CN | EC | FC | MA | NW | TB | TS | VS |
|---|---|---|---|---|---|---|---|---|---|---|---|---|---|---|---|---|---|---|---|---|---|---|---|---|
| BK | | 0 | 0 | 0 | 0 | 0 | 0 | 0 | 0 | 0 | 0 | 0 | 0 | 0 | 0 | 0 | 0 | 0 | 0 | 0 | 0 | 0 | 0 | 0 |
| MK | 0.45 | | 0 | 0 | 0 | 0 | 0 | 0 | 0 | 0 | 0 | 0 | 0 | 0 | 0 | 0 | 0 | 0 | 0 | 0 | 0 | 0 | 0 | 0 |
| TB | 0.67 | 0.59 | | 0 | 0 | 0 | 0 | 0 | 0 | 0 | 0 | 0 | 0 | 0 | 0 | 0 | 0 | 0 | 0 | 0 | 0 | 0 | 0 | 0 |
| BL | 0.67 | 0.55 | 0.72 | | 0 | 0 | 0 | 0 | 0 | 0 | 0 | 0 | 0 | 0 | 0 | 0 | 0 | 0 | 0 | 0 | 0 | 0 | 0 | 0 |
| CO | 0.33 | 0.34 | 0.6 | 0.57 | | 0 | 0 | 0 | 0 | 0 | 0 | 0 | 0 | 0 | 0 | 0 | 0 | 0 | 0 | 0 | 0 | 0 | 0 | 0 |
| CB | 0.76 | 0.7 | 0.83 | 0.96 | **0.73** | | 0 | 0 | 0 | 0 | 0 | 0 | 0 | 0 | 0 | 0 | 0.8 | 0 | 0 | 0 | 0 | 0 | 0 | 0 |
| KR | 0.66 | 0.55 | 0.7 | 0.97 | 0.56 | 0.95 | | 0 | 0.77 | 0 | 0 | 0 | 0 | 0 | 0 | 0 | 0 | 0 | 0 | 0 | 0 | 0 | 0 | 0 |
| B4 | 0.65 | 0.52 | 0.69 | 0.92 | 0.54 | 0.95 | 0.91 | | 0 | 0 | 0 | 0 | 0 | 0 | 0 | 0 | 0 | 0 | 0 | 0 | 0 | 0 | 0 | 0 |
| KB | 0.67 | 0.55 | 0.7 | 0.96 | 0.57 | 0.95 | 0 | 0.91 | | 0 | 0 | 0 | 0 | 0 | 0 | 0 | 0 | 0 | 0 | 0 | 0 | 0 | 0 | 0 |
| B1 | 0.66 | 0.54 | 0.71 | 0.97 | 0.56 | 0.96 | 0.97 | 0.84 | 0.97 | | 0 | 0 | 0 | 0 | 0 | 0 | 0 | 0 | 0 | 0 | 0 | 0 | 0 | 0 |
| NN | 0.31 | 0.44 | 0.71 | 0.75 | 0.35 | 0.9 | 0.75 | 0.71 | 0.75 | 0.75 | | 0 | 0 | 0 | 0 | 0 | 0 | 0 | 0 | 0 | 0 | 0 | 0 | 0 |
| VR | 0.64 | 0.55 | 0.62 | 0.73 | 0.56 | 0.85 | 0.68 | 0.69 | 0.69 | 0.72 | 0.69 | | 0 | 0 | 0 | 0 | 0 | 0 | 0 | 0 | 0 | 0 | 0 | 0 |
| EB | 0.67 | 0.56 | 0.69 | 0.97 | 0.58 | 0.96 | 0.97 | 0.91 | 0.97 | 0.97 | 0.75 | 0.72 | | 0 | 0 | 0.55 | 0 | 0 | 0 | 0 | 0 | 0 | 0 | 0 |
| MB | 0.39 | 0.06 | 0.56 | 0.51 | 0.28 | 0.68 | 0.5 | 0.47 | 0.5 | 0.5 | 0.37 | 0.52 | 0.52 | | 0 | 0 | 0 | 0 | 0 | 0 | 0 | 0 | 0 | 0 |
| MS | 0.5 | 0.19 | 0.68 | 0.68 | 0.36 | 0.89 | 0.68 | 0.63 | 0.68 | 0.67 | 0.52 | 0.64 | 0.7 | 0.16 | | 0 | 0 | 0 | 0 | 0 | 0 | 0 | 0 | 0 |
| EB | 0.67 | 0.56 | 0.69 | 0.97 | 0.58 | 0.96 | 0.97 | 0.91 | 0.97 | 0.97 | 0.75 | 0.72 | 0 | 0.52 | 0.7 | | 0 | 0 | 0 | 0 | 0 | 0 | 0 | 0 |
| CN | 0.7 | 0.6 | 0.76 | 0.97 | 0.62 | 0.01 | 0.96 | 0.94 | 0.96 | 0.97 | 0.82 | 0.75 | 0.97 | 0.57 | 0.78 | 0.97 | | 0 | 0 | 0 | 0 | 0 | 0 | 0 |
| EC | 0.66 | 0.49 | 0.79 | 0.96 | 0.56 | 0.96 | 0.96 | 0.94 | 0.96 | 0.96 | 0.79 | 0.81 | 0.97 | 0.45 | 0.67 | 0.97 | 0.97 | | 0 | 0 | 0 | 0 | 0 | 0 |
| FC | 0.68 | 0.58 | 0.72 | 0.96 | 0.59 | 0.96 | 0.95 | 0.92 | 0.95 | 0.97 | 0.78 | 0.72 | 0.96 | 0.54 | 0.73 | 0.96 | 0.96 | 0.96 | | 0 | 0 | 0 | 0 | 0 |
| MA | 0.65 | 0.55 | 0.66 | 0.83 | 0.56 | 0.92 | 0.69 | 0.79 | 0.69 | 0.82 | 0.72 | 0.6 | 0.83 | 0.51 | 0.66 | 0.83 | 0.87 | 0.89 | 0.83 | | 0 | 0 | 0 | 0 |
| NW | 0.31 | 0.4 | 0.7 | 0.75 | 0.31 | 0.9 | 0.74 | 0.7 | 0.74 | 0.75 | 0.24 | 0.67 | 0.75 | 0.33 | 0.47 | 0.75 | 0.82 | 0.79 | 0.78 | 0.7 | | 0 | 0 | 0 |
| TB | 0.67 | 0.55 | 0.65 | 0.93 | 0.56 | 0.96 | 0.93 | 0.87 | 0.93 | 0.93 | 0.74 | 0.73 | 0.86 | 0.51 | 0.67 | 0.87 | 0.95 | 0.95 | 0.94 | 0.81 | 0.73 | | 0 | 0 |
| TS | 0.7 | 0.6 | 0.7 | 0.9 | 0.6 | 0.9 | 0.9 | 0.9 | 0.9 | 0.9 | 0.8 | 0.7 | 0.9 | 0.5 | 0.7 | 0.9 | 0.9 | 0.9 | 0.9 | 0.8 | 0.8 | 0.5 | | 0 |
| VS | 0.64 | 0.54 | 0.63 | 0.79 | 0.55 | 0.9 | 0.76 | 0.75 | 0.76 | 0.78 | 0.7 | 0.45 | 0.79 | 0.5 | 0.64 | 0.79 | 0.83 | 0.86 | 0.79 | 0.65 | 0.68 | 0.77 | 0.85 | |

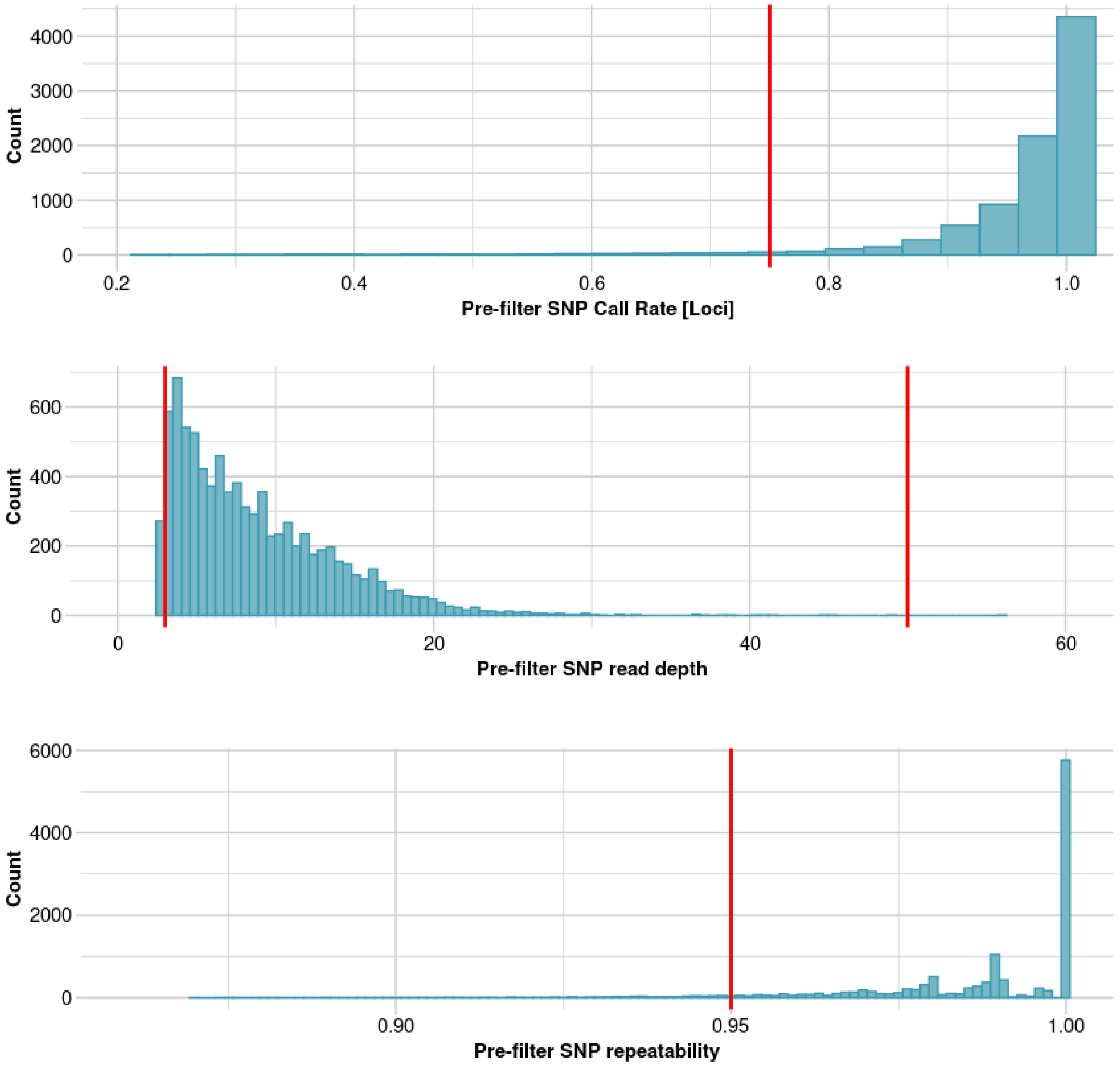


Fig. S4: Pre-filtering DaRT data sequence quality for a) call rate, b) read depth, and c) repeatability. Red lines indicate the filter threshold.

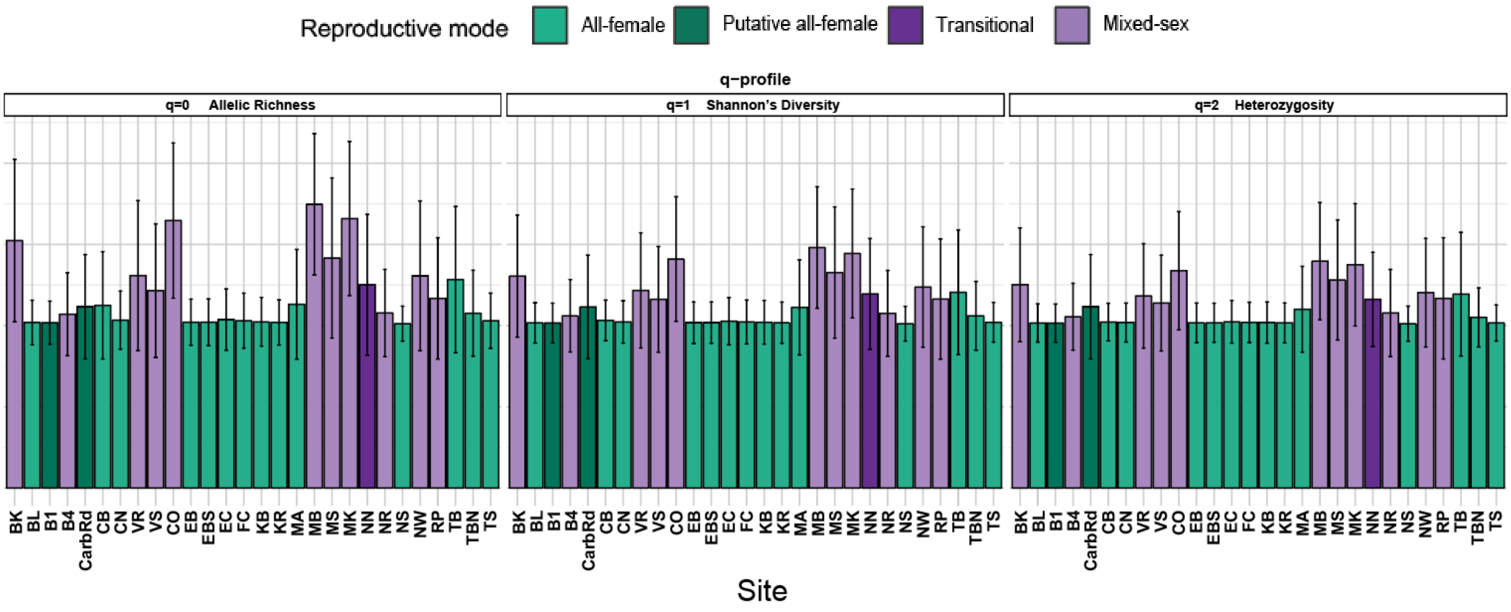


Fig. S5: Genetic diversity statistics for *M. batesii* individuals at each sampling location. Output of *gl.get.diversity()* showing various genetic diversity indices with colors representing site locations. Allelic richness ($^0H$) is the number of allelic types -1. Shannon's diversity ($^1H$) is an abundance-sensitive measure of species diversity with higher $^1H$ meaning there is higher uncertainty about what type to expect when randomly sampling one single allele. Heterozygosity ($^2H$) is the chance of choosing two different allelic types from the population. This measure of heterozygosity is distinct from the observed heterozygosity reported in the main text. The Y-axis is a common scale of effective numbers ($^qD$) that represent the number of equally frequent alleles that would be needed to yield the observed $^qH$ in the sample, which typically contains alleles at unequal frequencies (See Sherwin et al 2017). In all cases, all-female populations were less genetically diverse than mixed-sex populations.

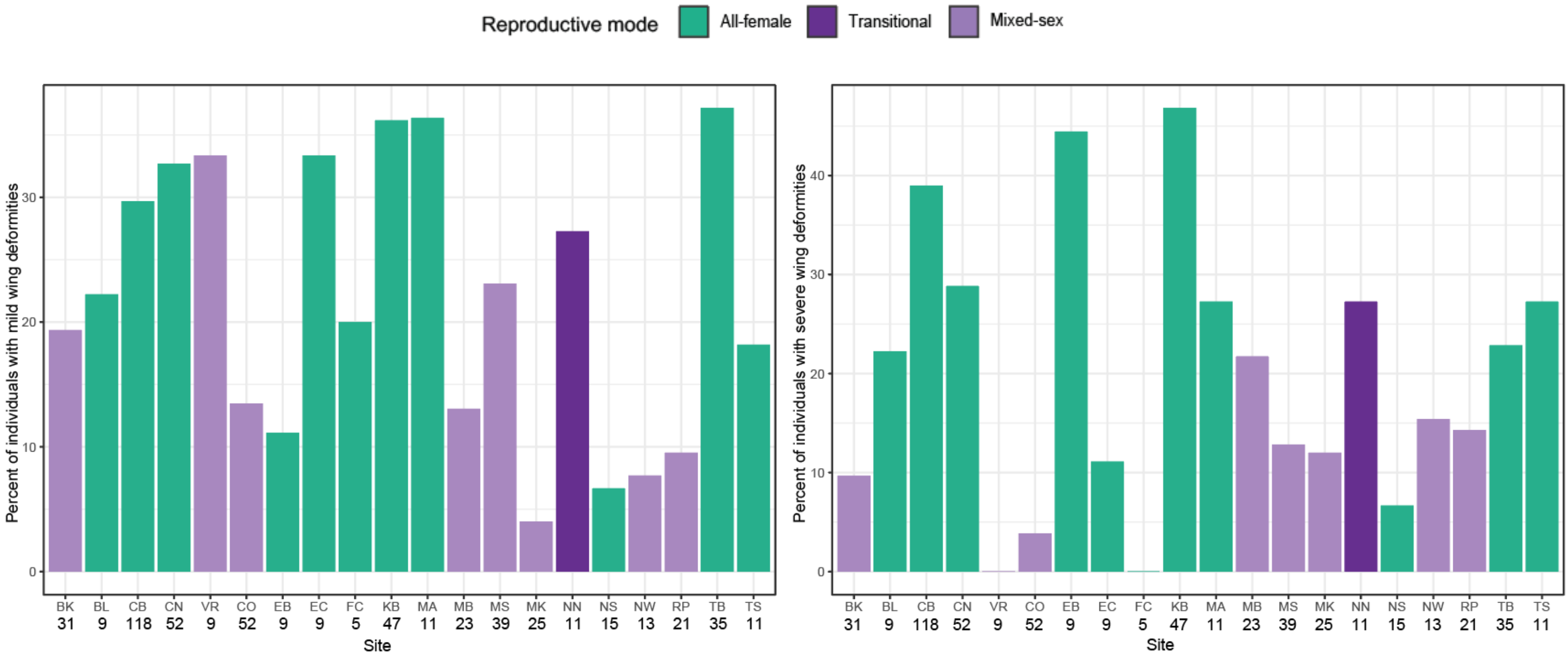


Fig. S6. Percentage of wild *M. batesii* individuals with mild and severe wing deformities at each of the sampling locations.

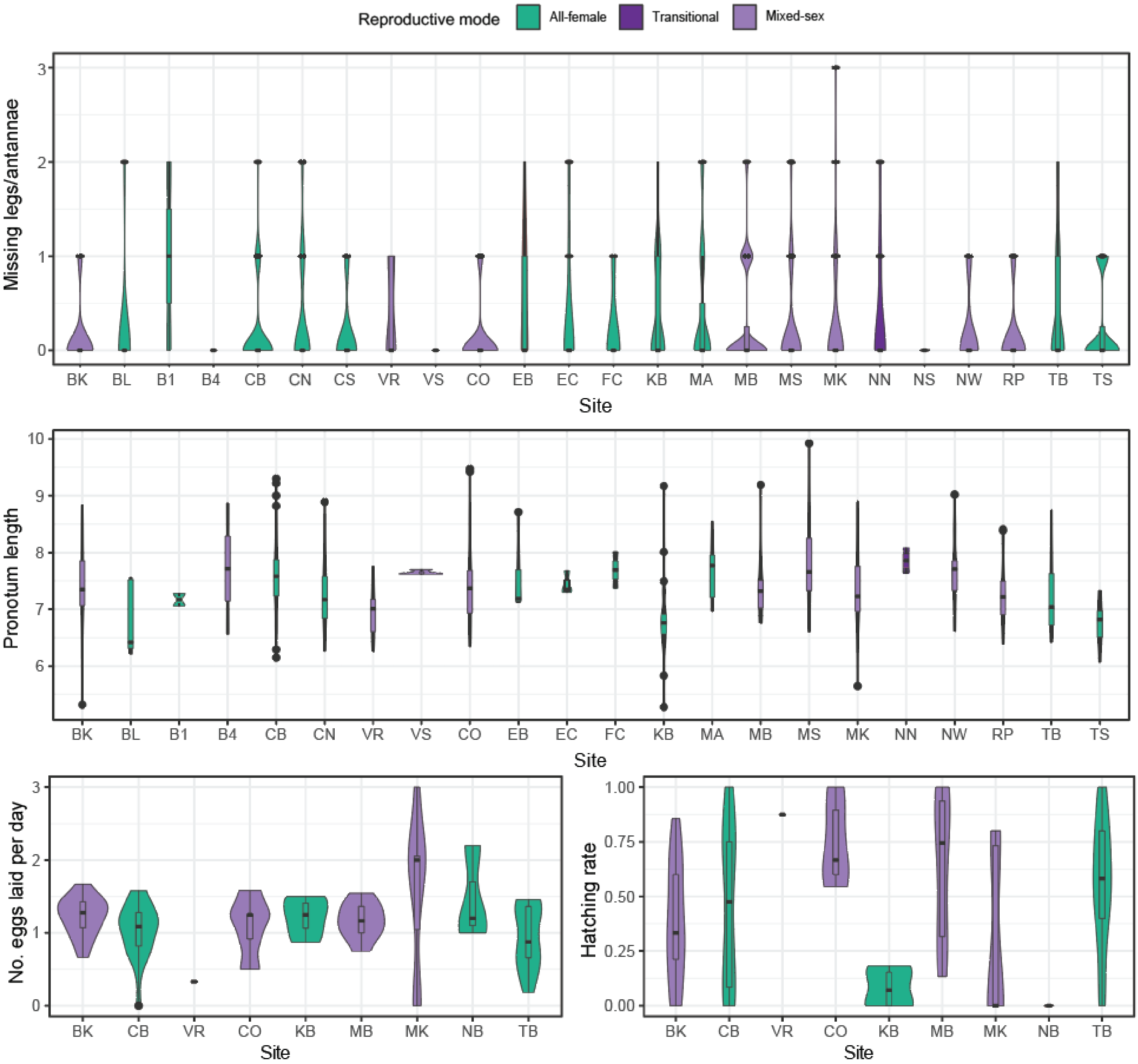


Fig. S7. Mean (bar), inter-quartile range (box) and non-outliner range (whisker) for phenotypic variables at *M. batesii* sampling locations. Violins indicate the distribution of the individual data at each location. Light purple fill indicates mixed-sex populations, Dark purple shows the transitional site NBN, dark green indicated populations that were putatively all-female (i.e., too few individuals were observed to determine sex ratio with confidence). Eggs were collected from a subset of the locations, providing data on fecundity (number of eggs laid per day) and hatching rate (proportion of eggs hatched).

Table S4: Hatchling sex ratios from *M. batesii* eggs collected in the field at several locations. Hatchling sex ratio in sites separated by year can be found in the data file "hatchling_sex_ratios_field_from_eggs.csv". Mixed-sex locations are shaded in purple.

| Site | Number of male hatchlings | Number of female hatchlings | Sex ratio (percent female) |
|---|---|---|---|
| BK | 49 | 43 | 0.467391 |
| BL | 0 | 25 | 1 |
| B1 | 0 | 20 | 1 |
| B4 | 1 | 2 | 0.666667 |
| CarbRd | 0 | 4 | 1 |
| CB | 0 | 179 | 1 |
| CN | 0 | 16 | 1 |
| VR | 4 | 7 | 0.636364 |
| VS | 10 | 8 | 0.444444 |
| CO | 25 | 17 | 0.404762 |
| EB | 0 | 1 | 1 |
| EC | 0 | 3 | 1 |
| KB | 0 | 4 | 1 |
| KR | 0 | 3 | 1 |
| Mad | 0 | 17 | 1 |
| MA | 0 | 5 | 1 |
| MB | 11 | 9 | 0.45 |
| MS | 5 | 6 | 0.545455 |
| MK | 48 | 43 | 0.472527 |
| NN | 16 | 18 | 0.529412 |
| NS | 0 | 10 | 1 |
| NW | 1 | 0 | 0 |
| NR | 13 | 11 | 0.458333 |
| RP | 0 | 1 | 1 |
| TB | 0 | 75 | 1 |
| TS | 0 | 9 | 1 |

Table S5: Filtering Steps used to remove low quality SNP calls and loci.

| Steps | Retained SNP count |
|---|---|
| Initial number of SNPs | 12977 |
| Polymorphic | 12518 |
| Reproducibility >= 0.95 | 11868 |
| Minor allele frequency >= 0.05 | 8999 |
| Call rate >= 75% | 8640 |
| Read depth >= 3 | 8367 |
| Hamming distance <= 0.8 | 7374 |
| Retained SNPs | 7374 |

Table S6: Sample sizes and summary statistics for the population genetics analyses. Sites with less than 2 *M. batesii* individuals sequenced were excluded from these analyses. nLoc is the number of loci within a population where there was information for at least one individual (out of 7374 loci total). Ho and HoSD are the observed heterozygosity of the population and the standard deviation, respectively. He and HeSD are the expected heterozygosity of the population under Hardy Weinberg equilibrium and the standard deviation, respectively. $F_{is}$ is the inbreeding coefficient.

| Site | Population type | Individuals sequenced | | Ho | HoSD | He | HeSD | $F_{is}$ |
|---|---|---|---|---|---|---|---|---|
| BK | Mixed-sex | 20 | 7374 | 0.131 | 0.183 | 0.149 | 0.188 | 0.142 |
| BL | All-female | 2 | 7303 | 0.015 | 0.114 | 0.009 | 0.062 | -0.323 |
| B1 | Putative all-female | 2 | 7291 | 0.015 | 0.115 | 0.008 | 0.061 | -0.355 |
| B4 | Mixed-sex | 2 | 7323 | 0.032 | 0.146 | 0.030 | 0.111 | 0.209 |
| CB | All-female | 15 | 7350 | 0.020 | 0.107 | 0.015 | 0.061 | -0.341 |
| CN | All-female | 4 | 7343 | 0.018 | 0.116 | 0.011 | 0.065 | -0.403 |
| VR | Mixed-sex | 7 | 7358 | 0.096 | 0.193 | 0.107 | 0.176 | 0.168 |
| VS | Mixed-sex | 4 | 7344 | 0.070 | 0.176 | 0.081 | 0.162 | 0.240 |
| CO | Mixed-sex | 13 | 7374 | 0.166 | 0.186 | 0.201 | 0.193 | 0.205 |
| EB | All-female | 2 | 7305 | 0.016 | 0.118 | 0.009 | 0.064 | -0.317 |
| EBS | All-female | 2 | 7301 | 0.016 | 0.116 | 0.009 | 0.064 | -0.288 |
| EC | All-female | 4 | 7321 | 0.016 | 0.111 | 0.014 | 0.072 | -0.011 |
| FC | All-female | 3 | 7296 | 0.018 | 0.116 | 0.011 | 0.067 | -0.311 |
| KB | All-female | 2 | 7319 | 0.018 | 0.123 | 0.010 | 0.067 | -0.311 |
| KR | All-female | 2 | 7279 | 0.016 | 0.118 | 0.009 | 0.064 | -0.310 |
| MA | All-female | 3 | 7345 | 0.017 | 0.112 | 0.056 | 0.146 | 0.741 |
| MB | Mixed-sex | 16 | 7374 | 0.151 | 0.156 | 0.236 | 0.188 | 0.381 |
| MS | Mixed-sex | 3 | 7372 | 0.174 | 0.267 | 0.162 | 0.201 | 0.106 |
| MK | Mixed-sex | 18 | 7374 | 0.183 | 0.195 | 0.219 | 0.199 | 0.186 |
| NN | Transitional | 4 | 7360 | 0.013 | 0.091 | 0.097 | 0.170 | 0.883 |
| NR | Mixed-sex | 1 | | | | | | |
| NS | All-female | 1 | | | | | | |
| NW | Mixed-sex | 3 | 7360 | 0.113 | 0.226 | 0.118 | 0.185 | 0.209 |
| RP | Mixed-sex | 1 | | | | | | |
| TB | All-female | 14 | 7369 | 0.021 | 0.096 | 0.101 | 0.192 | 0.796 |
| TBN | All-female | 2 | 7321 | 0.044 | 0.163 | 0.030 | 0.106 | -0.099 |
| TS | All-female | 4 | 7303 | 0.015 | 0.103 | 0.010 | 0.061 | -0.301 |

Table S7: MCMCglmm Model Specifications.

| | Iterations | Thinning interval | Burn in | MCMC sample size | DIC | Family |
|---|---|---|---|---|---|---|
| **Pronotum Length** | 3000000 | 1500 | 15000 | 1990 | 943.85 | Gaussian |
| **Fungus/Parasites** | 1000000 | 500 | 10000 | 1980 | 879.515 | Categorical (binomial) |
| **Missing Antennae/Legs** | 5000000 | 3000 | 40000 | 1654 | 851.77 | Poisson |
| **Mild Wing Deformities** | 5000000 | 1500 | 30000 | 3314 | 10.08 | Categorical (binomial) |
| **Severe Wing Deformities** | 5000000 | 1500 | 30000 | 3314 | 10.62 | Categorical (binomial) |
| **Fecundity (Eggs/day)** | 3000000 | 1000 | 10000 | 2990 | 196.84 | Poisson |
| **Egg Hatching Success** | 5000000 | 1500 | 50000 | 3300 | 1167.98 | Categorical (binomial) |

Table S8: Post-hoc Tukey test results.

| | diff | lwr | upr | p adj |
|---|---|---|---|---|
| swamp:asexual-beach:asexual | 18.05573 | -7.81712 | 43.92858 | 0.23473 |
| beach:sexual-beach:asexual | 5.82351 | -23.5135 | 35.16056 | 0.942291 |
| swamp:sexual-beach:asexual | -12.4464 | -41.7835 | 16.89061 | 0.635206 |
| beach:sexual-swamp:asexual | -12.2322 | -43.1562 | 18.69175 | 0.68349 |
| swamp:sexual-swamp:asexual | -30.5022 | -61.4261 | 0.421802 | 0.053966 |
| swamp:sexual-beach:sexual | -18.2699 | -52.1455 | 15.60556 | 0.444365 |
